# PRISM: A Plasmid-based Reporter for Intracellular Spectral Microscopy

**DOI:** 10.64898/2026.09.01.746207

**Authors:** FM Love, Z Baskir, T Allen, S Thomas, H Coyle, N Al-Dam, S Williams, H Maib, N Irigoyen, J Nixon-Abell

## Abstract

Organelles form an interconnected network whose morphology, positioning and interactions reflect cellular state. However, reproducibly quantifying these organelle phenotypes across large cell populations and diverse cell types remains a significant challenge. Here we present PRISM (Plasmid-based Reporter for Intracellular Spectral Microscopy), a PiggyBac-integrable construct encoding five unique fluorescent organelle reporters for spectral microscopy, with an accompanying modular analysis pipeline. PRISM stably labels the Golgi, peroxisomes, endoplasmic reticulum, mitochondria and lysosomes in multiple cell types while remaining compatible with additional molecular or functional probes. The workflow extracts over 500 metrics per cell, describing organelle morphology and distribution alongside pairwise and higher-order contacts. We use PRISM to characterise organelle responses to cytoskeletal perturbation, map PI(4)P redistribution during lysosomal damage, and reveal how Zika virus remodels the organelle landscape during infection. PRISM provides a reproducible approach for investigating organelle network remodelling across biological contexts.

## Introduction

Eukaryotic cells are characterised by their compartmentalisation into distinct membranebound organelles, but these compartments are neither static nor independent. Organelles form a dynamic network, and their abundance, size, shape, position, and interactions are continually remodelled in response to both cell-intrinsic and extracellular cues. This coordination allows fundamental cellular processes — signalling, metabolism, secretion — to be distributed across specialised organelles, and rapidly adjusted in response to such cues^1–3^. As a result, we can infer the functional state of a cell from the morphology and relationships between its constituent organelles. Recent studies have demonstrated that network-level organelle phenotypes can reliably distinguish different cell types, capture cellular responses to stress, and characterise the complex state transitions associated with differentiation^4–6^. However, challenges in labelling, imaging, and quantifying multiple types of organelles simultaneously, particularly in live cells, have restricted the broad adoption of network-level organelle phenotyping and limited its potential to deliver novel biological discoveries.

Spectral fluorescence microscopy has emerged as a key tool for addressing these challenges. While conventional fluorescence microscopy relies on selecting fluorophores with minimally overlapping emission spectra, spectral microscopy captures wavelength information along with intensity for each pixel, which allows multiple overlapping fluorophores to be imaged simultaneously and computationally unmixed^7^. This makes it possible to visualise multiple organelles together in a single acquisition, preserving their spatial relationships and wider network organisation. The major barrier to implementing spectral imaging at scale is maintaining reproducibility while labelling and quantifying multiple organelles across large cell populations, experimental conditions, and biological models. Existing approaches often require labelling cells with multiple independently-introduced reporters, which can limit throughput, introduce variability, and compromise downstream analysis.

Here, we present PRISM — a Plasmid-based Reporter for Intracellular Spectral Microscopy — which enables reproducible phenotyping of five organelles alongside additional molecular or functional markers in the same cells. PRISM encodes optimised fluorescent reporters for the Golgi apparatus, peroxisomes, endoplasmic reticulum (ER), mitochondria, and lysosomes within a single construct that can be integrated using the PiggyBac transposon system to generate stable lines in multiple cell types for spectral microscopy experiments. The fluorophore palette selected for PRISM leaves spectral capacity for additional labels, enabling the visualisation of specific proteins, lipids or functional reporters in the context of the wider organelle network. To accompany this, we have developed a flexible, modular image analysis workflow to extract quantitative metrics of organelle morphology and interactions, including both pairwise and higherorder organelle contacts from PRISM-labelled organelles as well as additional markers. We apply PRISM to investigate how the organelle network is remodelled in response to cytoskeletal perturbations, lysosomal damage, and Zika virus infection, and show that in each case it recovers known phenotypes while revealing previously unrecognised changes across the organelle network.

## Results

### PRISM: a polycistronic integration plasmid for labelling subcellular organelles across cell types

A major bottleneck to systematic organelle phenotyping is achieving consistent labelling of the same compartments across large cell populations and experimental series. To address this, we generated PRISM, a single polycistronic vector encoding fluorescent reporters for the Golgi apparatus, peroxisomes, endoplasmic reticulum, mitochondria and lysosomes (Fig. 1a). We selected these organelles because they are well characterised and present in most mammalian cell types. Together, they participate in functionally interconnected biosynthetic, metabolic and degradative pathways, and their distinct morphologies and distributions enable organelle remodelling to be examined across structurally and functionally diverse compartments. We selected organelle targeting sequences and fluorescent proteins to ensure clean and reliable localisation while minimising impact on organelle function or morphology (see Supplementary Text). Although the emission spectra of the five fluorophores overlap substantially, they are sufficiently distinct to be resolved by spectral imaging and linear unmixing (Fig. 1b). Their combined spectral range also leaves the blue (*<*450 nm) and far-red (*>*630 nm) regions free for additional user-selected markers.

**Figure 1.**
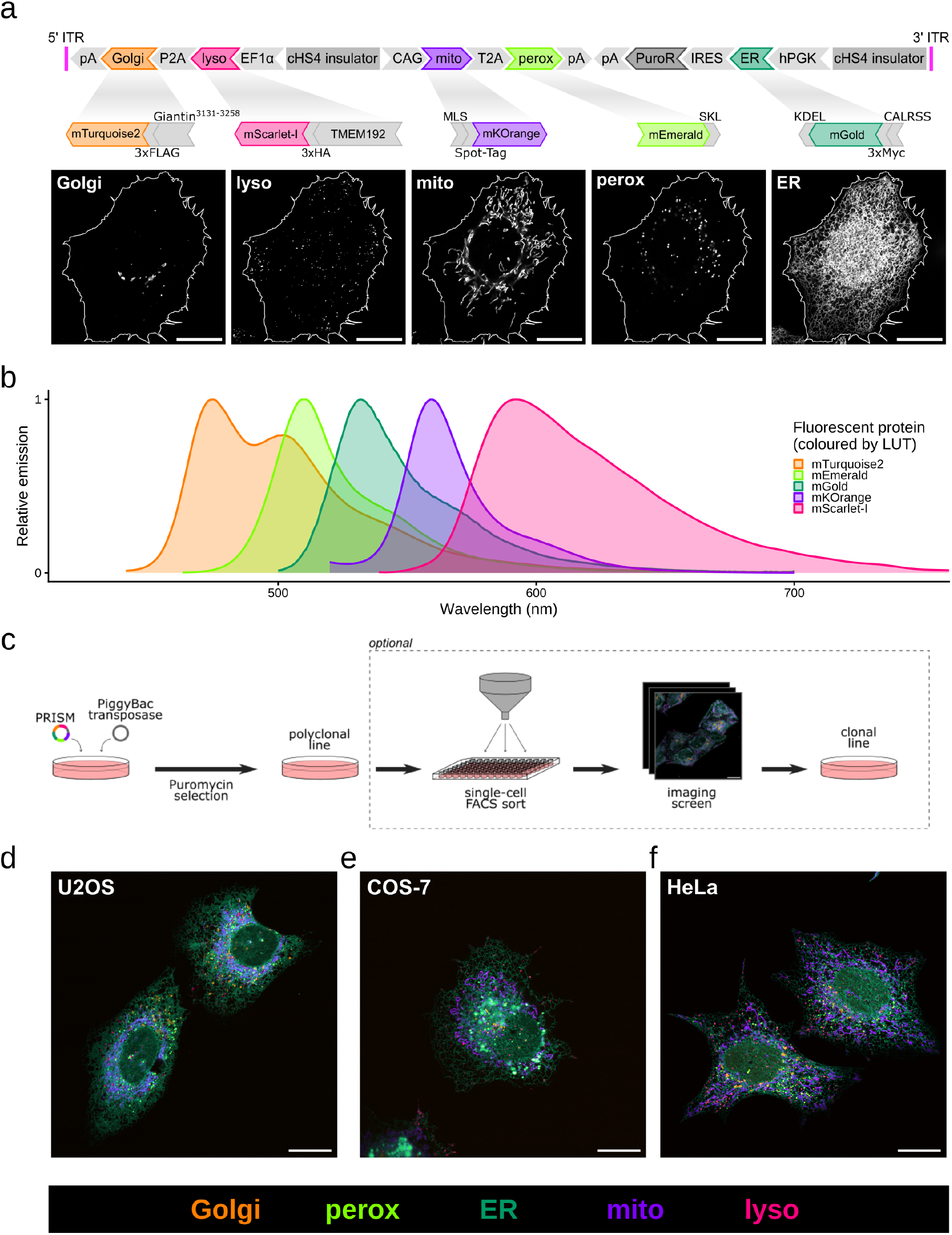
PRISM enables stable integration and spectrally resolved imaging of five organelles across multiple cell types. **(a)** Linear map of the PRISM construct between the PiggyBac inverted terminal repeats (ITRs), showing encoded reporters for the Golgi, lysosomes (lyso), mitochondria (mito), peroxisomes (perox) and endoplasmic reticulum (ER) together with a puromycin-resistance cassette. Each reporter comprises an organelle targeting sequence fused to a fluorescent protein, and, where indicated, a small epitope tag. Representative images of each reporter derived from a spectrally unmixed image of a cell expressing the complete PRISM construct are shown below. **(b)** Theoretical emission spectra of the fluorescent proteins used in PRISM, coloured according to the lookup tables (LUTs) used throughout this study. Corresponding excitation spectra are shown in Supplementary Fig. 1. **(c)** Schematic of stable PRISM integration. Co-transfection of the PRISM construct with PiggyBac transposase, followed by puromycin selection, generates a stable polyclonal population. Optional clonal selection can be carried out to obtain a more homogeneous population of cells. **(d-f)** Representative composite images of stable PRISM-expressing live **(d)** U2OS, **(e)** COS-7, and **(f)** clonal HeLa cells. Lookup tables for each organelle channel are indicated below. Individual channels are shown in Supplementary Fig. 2. All scale bars, 20 µm. Images shown are representative of cells from 3 independent experiments.

The various organelle reporters are flanked by PiggyBac inverted terminal repeats (ITRs), which enable stable genomic integration when PRISM is co-transfected alongside the PiggyBac transposase, as well as a puromycin resistance gene to select for successful integration (Fig. 1c). We used this system to generate stable PRISM-expressing populations in HeLa, U2OS and COS-7 cells. In each cell type, all five reporters were expressed and could be spectrally unmixed, with each reporter localising to the expected organelle (Fig. 1d-f, Supplementary Fig. 2). Where initial polyclonal populations show variable expression of the PRISM markers, clonal selection can be carried out to produce a more uniform line, which can improve spectral unmixing. We used FACS-based clonal isolation on the PRISM HeLa line, and the resultant clone was used for all experiments described in this paper (Individual channels in Supplementary Fig. 2).

**Figure 2.**
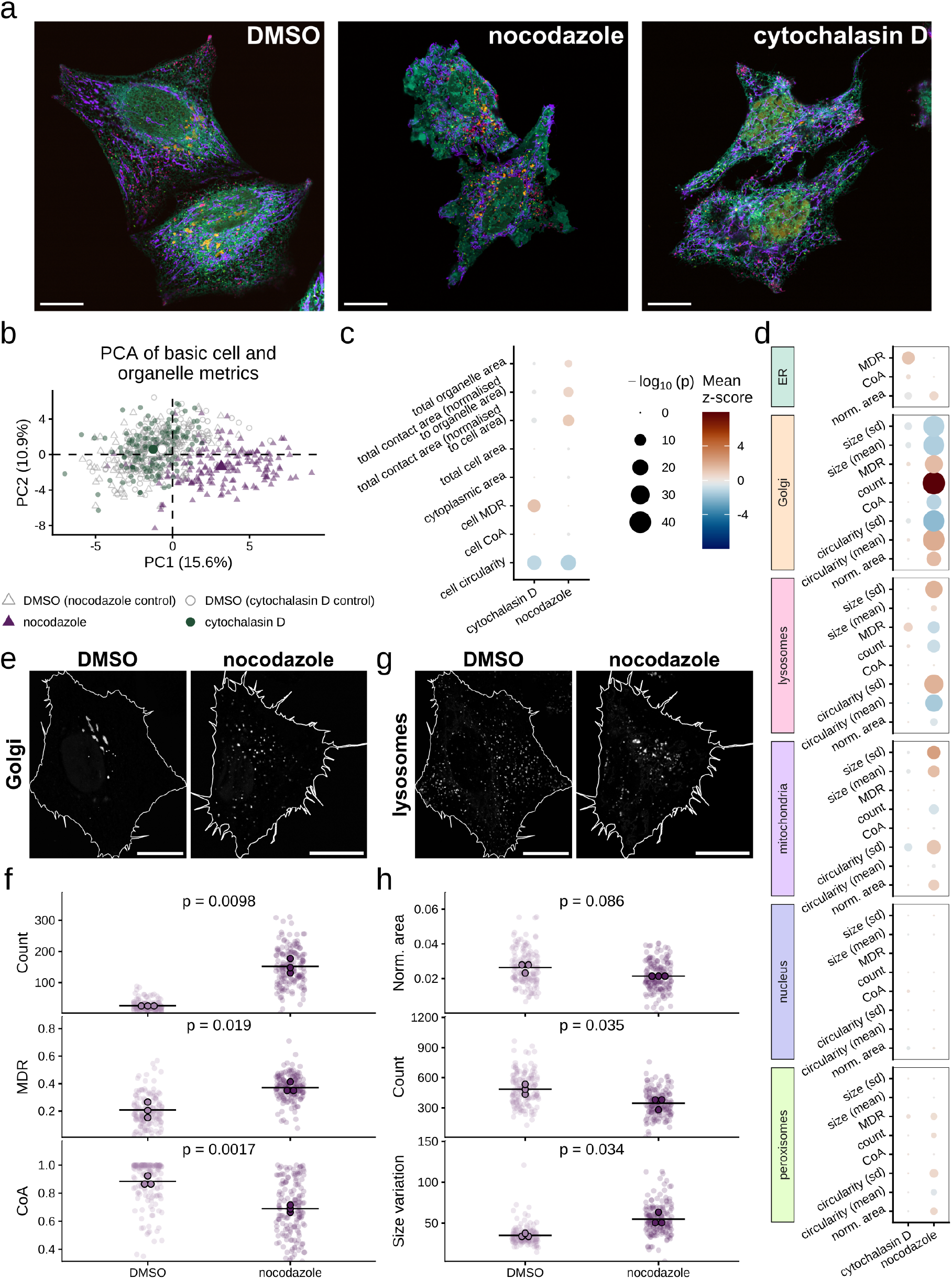
PRISM resolves established and previously unrecognised organelle responses to cytoskeletal disruption. **(a)** Representative images of live PRISM-expressing HeLa cells treated with DMSO, 5 µM nocodazole or 1 µM cytochalasin D for 1 hour. **(b)** PCA of the 50 standardised metrics presented in panels and (d). Each point represents an individual cell. n = 144 (DMSO nocodazole control), 157 (nocodazole), 150 (DMSO cytochalasin D control), and 154 (cytochalasin D) cells across 3 independent experiments. **(c)** Dot plot of cell-wide summary metrics in nocodazole- and cytochalasin D-treated cells relative to their respective DMSO controls. Point size indicates the adjusted *p*-value from a Wilcoxon rank-sum test with Benjamini-Yekutieli correction for multiple comparisons, colour indicates the mean z-score for each metric relative to the respective control. n numbers provided in (b). **(d)** Dot plot of organelle metrics. Scales for adjusted *p*-value and z-score are as defined in (c). **(e)** Representative images showing Golgi morphology in DMSO control and nocodazole-treated live cells. **(f)** Plots of the number of segmented Golgi objects (count), mean distribution radius (MDR), and coefficient of asymmetry (CoA) in DMSO control vs nocodazole-treated cells. Mean = black line. *P*-values from Welch’s t-test on independent replicate means, n = 3. Individual cell values are shown for reference, but not included in the statistics. **(g)** Representative images showing lysosome morphology in DMSO control and nocodazole-treated live cells. **(h)** Plots of normalised total lysosome area, the number of segmented lysosome objects (count), and variation in lysosome object size (s.d.) in DMSO control vs nocodazole-treated cells. Mean = black line. *P*-values from Welch’s t-test (two-sided, unpaired) on independent replicate means, n = 3. Individual cell values are shown for reference, but not included in the statistics. All scale bars, 20 µm. Images shown are representative of cells from 3 independent experiments. Details on all normalisations are provided in Methods.

### A modular analysis pipeline to quantify organelle phenotypes and network organisation

To convert PRISM images into quantitative organelle phenotypes, we developed a modular analysis pipeline to accompany the plasmid. While PRISM is suited for a range of 2D, 3D, live, and fixed imaging approaches, we have chosen to focus on 2D confocal imaging of single optical sections for this analysis, as a compromise between sampling breadth and imaging speed for adherent cells.

The pipeline consists of two custom ImageJ macros and an R Markdown notebook (Supplementary Fig. 3), which are run sequentially. The pipeline accepts cell, nuclear and organelle masks generated from the image data using standard segmentation tools such as ilastik, CellProfiler or Cellpose^8–10^. Users can select the organelles and additional labels to be measured, allowing the analysis to be tailored to each experiment without changing the underlying pipeline. From the segmentation masks, the pipeline extracts quantitative metrics describing organelle abundance, morphology and intracellular distribution. Distribution is described using mean distribution radius (MDR) and coefficient of asymmetry (CoA), which have been adapted from Zheng et al.^11^. MDR quantifies the radial position of an organelle between the nucleus and the cell periphery, ranging from 0 at the nucleus to 1 at the cell periphery. CoA quantifies the asymmetry of an organelle’s distribution around the nucleus, from 0 for a perfectly symmetrical distribution to 1 when the organelle lies entirely on one side of the nucleus. The pipeline also identifies pixel overlap between organelle masks, allowing pairwise and higherorder contacts involving three or more organelles to be quantified. At the spatial resolution used here, these regions encompass both direct membrane contacts and nearby membranes that cannot be spatially resolved from one another. We therefore refer to these measurements as ‘contacts’ throughout, rather than as *bona fide* membrane contact sites. Collectively, the basic PRISM analysis panel yields 507 metrics from each cell (see Supplementary Text). These measurements enable the systematic, quantitative assessment of organelle network organisation across hundreds of individual cells.

**Figure 3.**
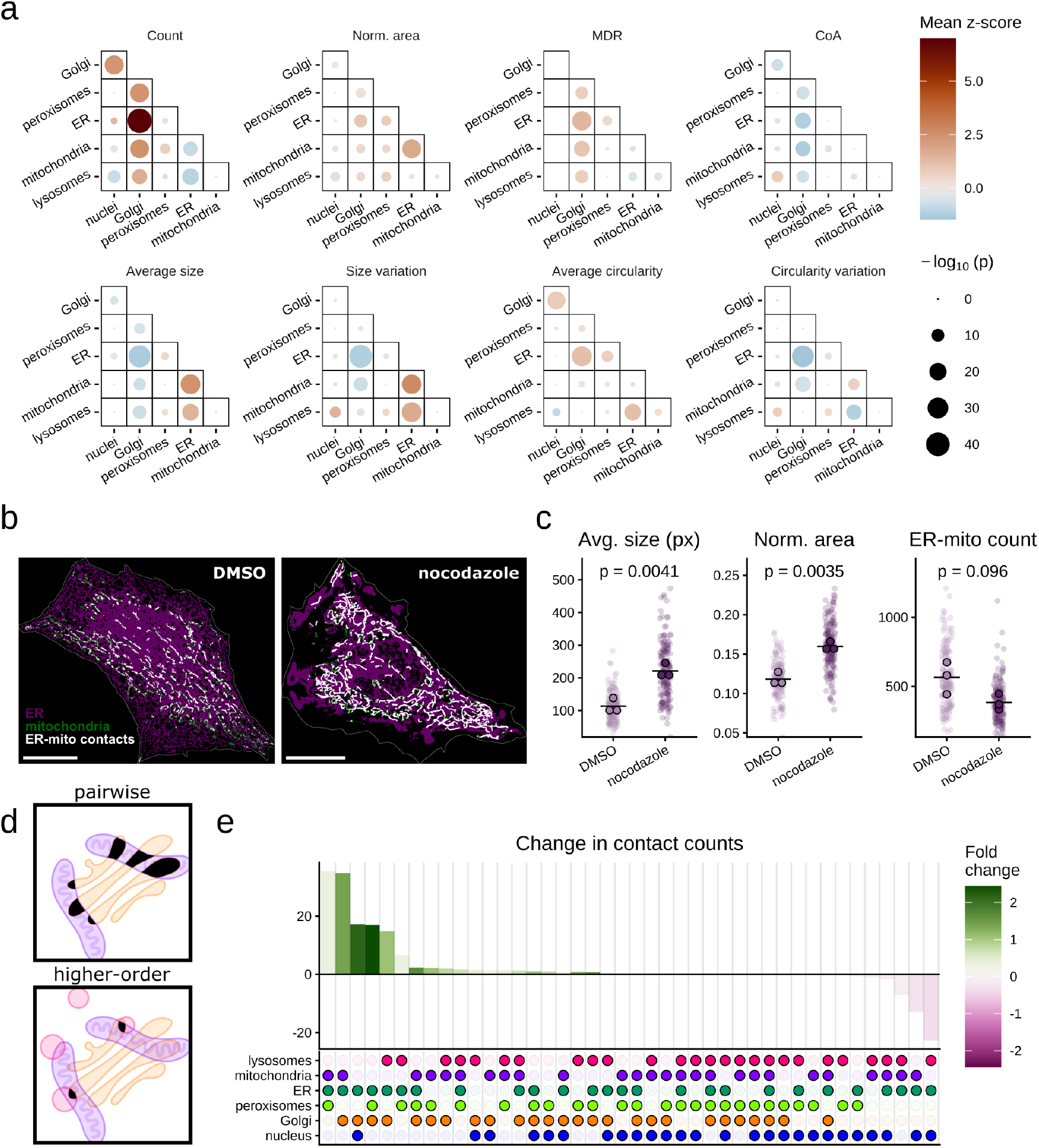
Microtubule depolymerisation remodels pairwise and higher-order contacts. **(a)** Dot plots of metrics describing the number (count), size, shape, and distribution of each pairwise organelle contact type imaged in DMSO control and nocodazole-treated cells. Point size indicates the adjusted *p*-value from a Wilcoxon rank-sum test with Benjamini-Yekutieli correction for multiple comparisons, colour indicates the mean z-score for each metric relative to the DMSO control. n = 144 (DMSO), 157 (nocodazole) cells across 3 independent experiments. **(b)** ER and mitochondria segmentation masks and the corresponding contact mask showing overlap between ER and mitochondria in representative DMSO control and nocodazole-treated live cells. Scale bar, 20 µm. **(c)** Plots showing the number (count), average size, and normalised total area of ER-mitochondrial contacts in nocodazole vs DMSO control cells. Mean = black line. *P*-values from Welch’s t-test (two-sided, unpaired) on independent replicate means, n = 3. Individual cell values are shown for reference but not included in the statistics. Schematic illustrating pairwise contacts involving two organelles and higher order contacts involving three or more organelles. **(e)** Bar plot showing the change in the mean number (count) of higher-order contacts between DMSO control and nocodazole-treated cells. The organelles involved in each contact are indicated by the dots below the X-axis. Bar height indicates the change in mean number of contacts, while the colour of each bar indicates the fold change relative to the DMSO control. Images shown are representative of cells from 3 independent experiments.

### PRISM reveals coordinated organelle remodelling following microtubule depolymerisation

To validate PRISM, we sought to expose cells to perturbations with known effects on cellular organisation to confirm that we could recover established phenotypes and determine whether we could identify additional changes to the organelle network. As such, we decided to investigate the effects of cytoskeletal disruption on organelle remodelling using nocodazole or cytochalasin D (Fig. 2a and Supplementary Fig. 4). Nocodazole prevents microtubule polymerisation, with documented effects on several organelles, most notably the Golgi, which becomes fragmented into distributed ‘ministacks’^12–14^. Cytochalasin D disrupts actin filaments and causes pronounced changes in cell shape^15^. The clonal PRISM HeLa line was treated with 5 µM nocodazole or 1 µM cytochalasin D, and images were processed through our analysis pipeline to extract organelle metrics.

**Figure 4.**
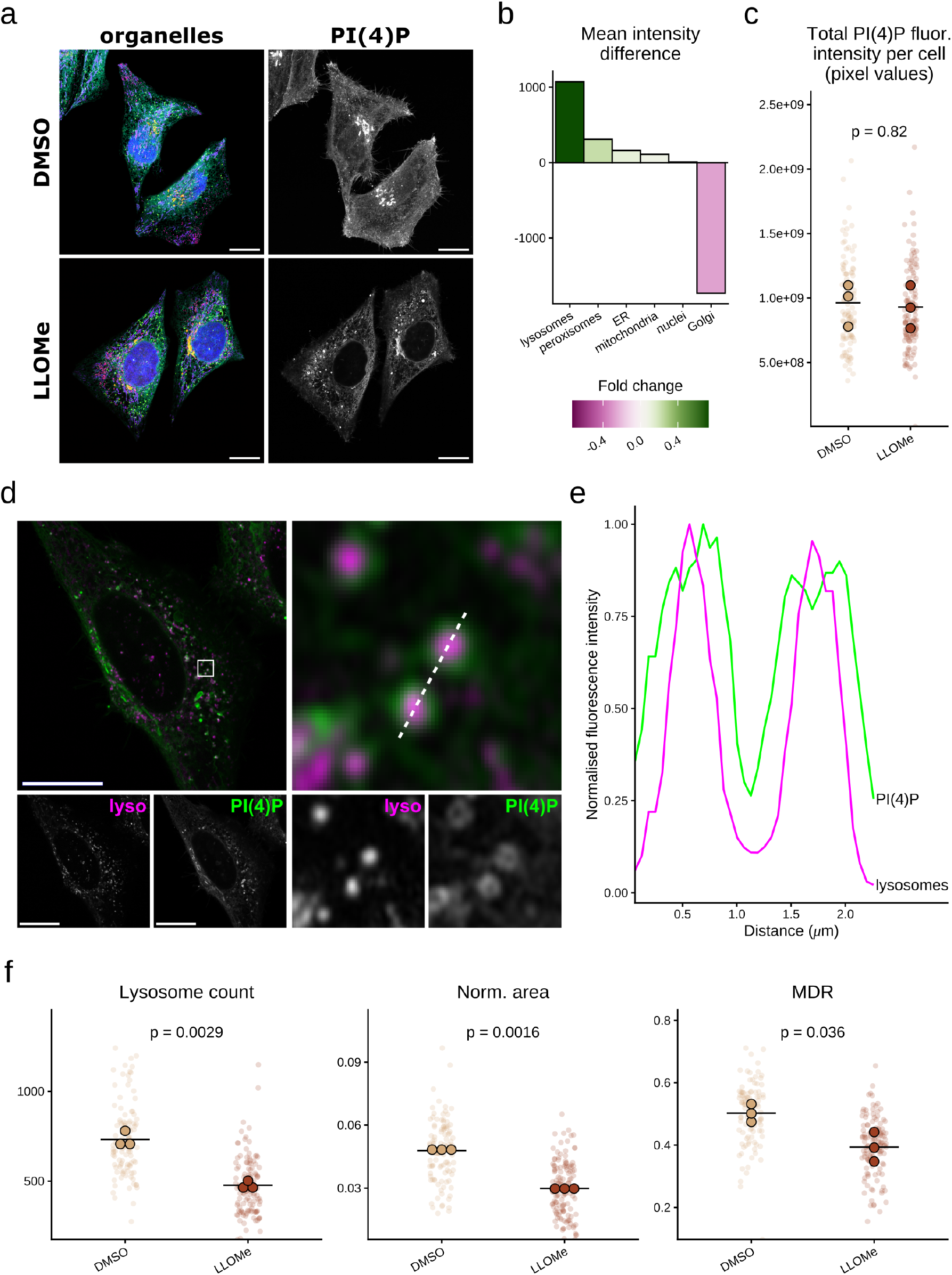
Lysosomal damage redistributes PI(4)P from the Golgi to juxta-lysosomal membranes. **(a)** Representative images of the PRISM HeLa line treated with DMSO or 0.5 mM LLOMe for 1 h, fixed, and co-stained with DAPI and an AF647-conjugated PI(4)P biosensor. Individual channels are shown in Supplementary Fig. 5. **(b)** Bar plot showing the change in mean PI(4)P fluorescence intensity across labelled organelles between DMSO control and LLOMe-treated cells. Bar height indicates the difference in mean fluorescence intensity, while bar colour indicates the fold change relative to the control. n = 108 (DMSO), 125 (LLOMe) cells across 3 independent experiments. **(c)** Plot showing the total PI(4)P fluorescence intensity (raw integrated density within the segmented cell ROI) in DMSO control and LLOMe-treated cells. Mean = black line. *P*-values from Welch’s t-test (two-sided, unpaired) on independent replicate means, n = 3. Individual cell values are shown for reference, but not included in the statistics. **(d)** Representative image of lysosome and PI(4)P channels from a LLOMe-treated cell. Right is a 4.12×4.12 µm zoom panel (white box in left image) showing PI(4)P-labelled structures surrounding lysosomes. **(e)** Intensity profile across the white line indicated in (d). Intensity values are normalised to the maximum value in each profile. **(f)** Plots showing the number (count) of segmented lysosomal objects, normalised total lysosomal area and mean distribution radius (MDR) in DMSO control and LLOMe-treated cells. Mean = black line. *P*-values from Welch’s t-test (two-sided, unpaired) on independent replicate means, n = 3. All scale bars, 20 µm. Images shown are representative of cells from 3 independent experiments.

We first used principal component analysis (PCA) to examine the global response across our cell-wide and organelle metrics (Fig. 2b). The two DMSO controls and cytochalasin D condition cluster together, while the nocodazole-treated population is clearly separated, indicating more substantial remodelling with this treatment.

We then separated the metrics by category to examine cell-wide and organelle measurements individually. Both nocodazole and cytochalasin D reduced cell circularity (mean z-scores −1.33 and −1.13, respectively), confirming that each treatment altered overall cell shape (Fig. 2c). Cytochalasin D-treated cells also showed an increase in cell MDR (mean z-score 1.24), indicating that a greater proportion of the cell area was distributed at larger radial distances from the nucleus, which reflects the irregular cell shapes caused by actin filament disruption. However, changes in the cell-wide organelle and contact metrics were far more pronounced following nocodazole treatment (Fig. 2c). This distinction became clearer when we examined the measurements for each organelle individually (Fig. 2d). Most organelle phenotypes were minimally affected by cytochalasin D treatment, whereas nocodazole substantially altered features across several compartments. The strongest responses were observed in Golgi and lysosomal metrics, with more modest changes across ER, mitochondria and peroxisomes. Although disruption of either cytoskeletal network dramatically affected cell morphology, widespread remodelling of the measured organelles was primarily associated with microtubule disruption. This is consistent with several previous studies that have identified microtubules as major determinants of organelle shape and positioning (reviewed in^16^).

As previously reported, nocodazole treatment dispersed the compact perinuclear Golgi into numerous fragments distributed throughout the cell (Fig. 2e). This was reflected by a substantial increase in the number of segmented Golgi objects. Additionally, the fragmented Golgi was positioned further from the nucleus, reflected by an increase in MDR, while the decrease in CoA illustrated that the Golgi fragments were more uniformly distributed around the nucleus (Fig. 2f). These measurements therefore reflect the established morphological and spatial features of nocodazole-induced Golgi fragmentation.

We also identified an unexpected lysosomal phenotype in response to nocodazole treatment, with lysosomes appearing less evenly dispersed and forming larger, irregular structures (Fig. 2g). Although total lysosomal area was not significantly altered, the number of segmented lysosomal objects decreased, and the variability in their size increased (Fig. 2h). The role of microtubule-motor transport in lysosome positioning is well-established^17, 18^, and we interpret these measurements to reflect lysosomal clumping following the disruption of these transport mechanisms.

### Microtubule disruption remodels pairwise and higher-order organelle contacts

Having established which organelle phenotypes are altered by nocodazole, we next asked how microtubule disruption affected contacts between compartments. We quantified all fifteen possible pairwise contacts (i.e. between two organelles) among the labelled organelles, comparing changes in contact number, area and intracellular distribution (Fig. 3a). Contacts involving the Golgi showed the most widespread changes, consistent with its extensive fragmentation and redistribution. However, several contacts involving the ER were also substantially altered, with a particularly pronounced response observed for ER-mitochondria contacts (Fig. 3b). Following nocodazole treatment, the average size of each contact increased, producing an overall increase in total contact area despite no significant change (although a downward trend) in the number of contacts (Fig. 3c). These data suggest that microtubule disruption might reorganise ER-mitochondria interfaces, although whether these changes reflect true alterations in ER-mitochondria membrane contact sites remains to be determined.

A key advantage of the PRISM system is the ability to look beyond pairs of organelles to interrogate higher-order contacts, where three, four, or more organelles might be interacting (Fig. 3d). Beyond spectral imaging, there are no other existing techniques that permit this type of analysis. For all combinations of labelled organelles, we calculated both the change in absolute contact number and the corresponding fold change in the nocodazole condition relative to the DMSO control (Fig. 3e). As expected, many of the strongest responses involved Golgi contacts, consistent with its remodelling following nocodazole treatment. However, ER-peroxisome-mitochondria and ER-peroxisome-lysosome contacts were also among the most strongly increased higher-order combinations (fold-changes of 0.37 and 0.33, respectively). Functional tripartite contacts involving these sets of organelles have been recently described^19–21^, but their sensitivity to microtubule disruption has yet to be explored. These data suggest that the organisation of these contacts depends on the microtubule network, and illustrate the utility of PRISM in identifying specific higher-order organelle relationships for further mechanistic investigation. Taken together, these analyses show that nocodazole-induced remodelling extends from individual organelle phenotypes to pairwise and higher-order relationships across the organelle network.

### PRISM maps PI(4)P redistribution during lysosomal damage

In addition to extracting organelle phenotypes, PRISM also enables quantification of additional molecular probes with respect to all five labelled organelles in the same cell. Here we demonstrate this by mapping changes in phosphatidylinositol 4-phosphate (PI(4)P) distribution in response to lysosomal damage, which prior work^22^ has suggested causes its accumulation on damaged lysosomes. To induce lysosomal damage, we treated PRISM HeLa cells with 0.5 mM LLOMe for 1 hour before fixing the cells and labelling with an Alexa Fluor 647-conjugated PI(4)P biosensor (Fig. 4a, individual channels in Supplementary Fig. 5).

**Figure 5.**
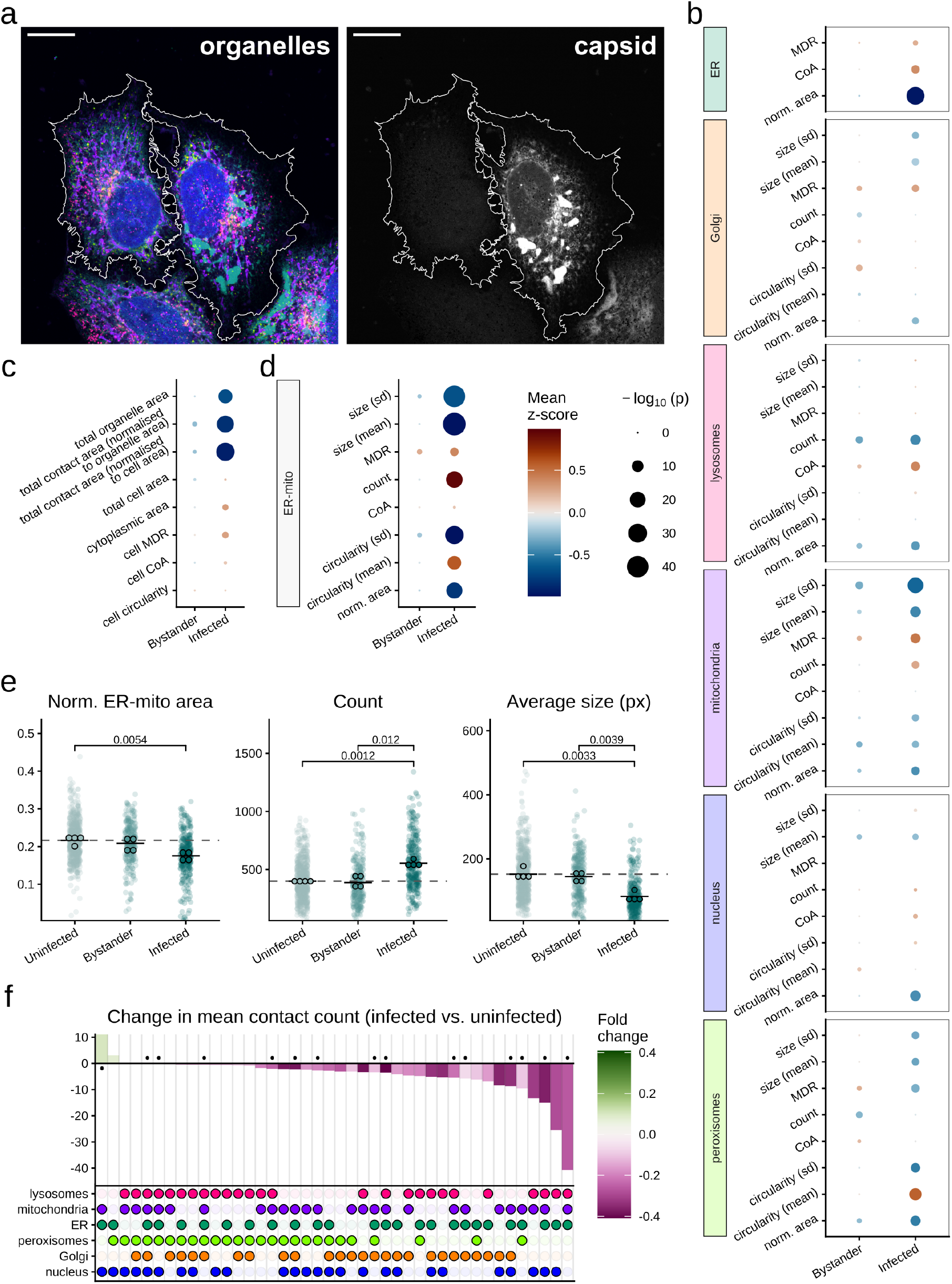
PRISM reveals coordinated changes in network organisation in response to ZIKV infection. **(a)** Representative image of the PRISM HeLa line infected with ZIKV encoding JFX646-Halo labelled capsid, fixed and co-stained with DAPI. The cell on the right is infected, while the cell on the left is an uninfected bystander. Scale bar, 20 µm. **(b)** Dot plot of aggregate organelle metrics for bystander and infected cells relative to an uninfected control. Point size indicates the adjusted *p*-value from a Wilcoxon rank-sum test with Benjamini-Yekutieli correction for multiple comparisons, colour indicates the mean z-score for each metric relative to the DMSO control. n = 263 (bystander), 289 (infected), 630 (uninfected) cells from 4 independent experiments. **(c)** Dot plot of cell-wide summary metrics. Scales for adjusted *p*-value and z-score, and n numbers are as defined in (b) **(d)** Dot plot of ER-mitochondrial contact metrics. Scales for adjusted *p*-value and z-score, and n numbers are as defined in (b). Data for all pairwise contact metrics shown in Supplementary Fig. 6. **(e)** Plots showing normalised total area, number (count), and average size of ER-mitochondria contacts in uninfected, bystander, and ZIKV infected conditions. Mean = solid black line, comparison to uninfected mean = dashed gray line. *P*-values are from pairwise Welch’s t-tests (two-sided, unpaired) with Bonferroni correction for multiple comparisons on independent replicate means, n = 4. All pairs compared; non-significant comparisons (*p >*0.05) are not shown. Individual cell values are shown for reference but not included in the statistics. **(f)** Bar plot showing the change in the mean number (count) of higher-order contacts between ZIKV infected and uninfected cells. The organelles involved in each contact are indicated by the dots below the X-axis. Bar height indicates the change in mean number of contacts, while the colour of each bar indicates the fold change relative to the uninfected control. Dots above or below each bar indicate higher-order contacts involving ER and mitochondria. Images shown are representative of cells from 4 independent experiments. Details on all normalisations are provided in Methods.

In DMSO-treated control cells, the most prominent PI(4)P pool was associated with the Golgi. Following treatment with LLOMe, we observed a substantial increase in PI(4)P colocalisation with lysosomes (fold-change 0.73), with a concomitant decrease in signal at the Golgi (fold-change −0.27) (Fig. 4b). Given that we see no change in total cell-wide PI(4)P levels with LLOMe treatment (Fig. 4c), these data suggest that PI(4)P is relocalised from the Golgi to lysosomes in response to lysosomal damage. Further experiments are necessary to determine whether PI(4)P is transported directly, or if the shift in localisation is due to transport of its kinase PI4K2A from the Golgi towards damaged lysosomes, as other studies have indicated^22, 23^.

It is also apparent from our imaging data that, at the 1 hour timepoint, much of the PI(4)P does not localise directly to lysosomal membranes; rather, it appears to label structures that closely associate with, and sometimes engulf, lysosomes (Fig. 4d). Line profiles across representative structures show PI(4)P intensity peaks flanking the lysosomal membrane signal (Fig. 4e). These structures are reminiscent of the autophagic membranes that enclose damaged lysosomes during lysophagy, and likely reflect PI(4)P’s established localisation to autophagic organelles^24^.

Consistent with this interpretation, LLOMe treatment reduced both the number and total area of segmented lysosomes and decreased lysosomal MDR, reflecting a shift towards a more perinuclear distribution (Fig. 4f). All of these signatures are suggestive of lysophagic clearance following LLOMe-induced damage. This is broadly in agreement with previous reporting —– while exposure to LLOMe initially triggers the rapid recruitment of ESCRT and membrane repair machinery, this localisation declines after ~30 minutes of exposure, and is followed by an increase in markers of lysophagy^25–27^.

Together, these data demonstrate the value of PRISM in generating unbiased organelle labelling and phenotyping. We recover the previously-reported association between PI(4)P and damaged lysosomes, while also revealing a decrease in Golgi localisation, and we can further link this to changes in lysosome abundance and positioning in the same cells.

### PRISM reveals network-wide organelle remodelling during ZIKV infection

Intracellular pathogens often extensively remodel host cell organelles — both to promote replication and to evade the immune response — while host cells themselves also alter their organelles in response to the infection^28, 29^. However, the pattern of these changes varies between pathogens and remains poorly mapped, limiting our understanding of how infection reshapes the organelle network and highlighting a significant research need. PRISM provides a valuable tool to interrogate the host-pathogen interaction by enabling the investigation multiple organelles simultaneously during infection, revealing wider patterns of network remodelling.

We applied PRISM to study Zika virus (ZIKV) infection in our clonal HeLa line. ZIKV has previously been shown to co-opt ER membranes into replication factories, fragment mitochondria, and lead to a loss of peroxisomes in infected cells^30–33^. We therefore sought to determine whether we could recover these established phenotypes and identify a broader reorganisation of the host cell organelle landscape.

To identify infected cells, we generated a ZIKV mutant virus encoding a HaloTag-fused capsid protein (Supplementary Fig. 6), which could be detected using the JFX646 far-red HaloTag ligand alongside the five organelle reporters (Fig. 5a, individual channels in Supplementary Fig. 6). This enabled us to distinguish infected cells from ‘bystander’ cells (uninfected cells on the same coverslip), allowing potential non-cell-autonomous responses to infection to be identified. Because the PRISM reporter was stably expressed throughout the population, organelle phenotypes could be measured in uninfected, bystander and infected cells in a uniform background. This is particularly valuable for infection models, in which only a subset of the exposed population might be infected by the pathogen.

We first examined the phenotype of each organelle individually (Fig. 5b). Infected cells showed a substantial reduction in the measured ER total area, consistent with the previously reported remodelling and compaction of ER membranes around ZIKV replication factories^30^. Mitochondria became smaller and more numerous, recovering the established mitochondrial fragmentation phenotype observed following ZIKV infection^31, 32^. We also detected a reduction in peroxisomal area, consistent with the reported depletion of peroxisomes from infected cells^33^ (Supplementary Fig. 7). In addition, we observed fewer significant changes in the bystander population and consistently smaller z-scores, reflecting less substantial organelle remodelling. The PRISM pipeline therefore recovered established ZIKV-associated phenotypes across several organelles within a single acquisition.

Comparing the cell-wide metrics, we also noted ZIKV infection caused a reduction in total organelle contact area (mean z-score −0.92) which remained pronounced after normalisation to total organelle area (mean z-score −0.84), indicating that it could not be explained solely by the accompanying decrease in segmented organelle area (Fig. 5c). A comparison of all pairwise organelle contacts (Supplementary Fig. 6) identified ER-mitochondria contacts as undergoing the most substantial change (Fig. 5d). While the total number of these contacts increases in the infected condition, each individual contact is smaller, and the net result is a decrease in total contact area (Fig. 5e). In support of these data, it has recently been shown that ZIKV infection decreases ER-mitochondria membrane contact sites through down-regulation of the RRPB1 tether protein^34^. The broad reduction in organelle contacts also extended to higher-order contacts (Fig. 5f). Most higher-order contacts were decreased in ZIKV-infected cells, contrasting with the smaller and more balanced changes observed following nocodazole treatment (Fig. 3e). The largest reduction involved ER-mitochondria-lysosome contacts, which have been shown to play a role in establishing mitochondrial morphology and lipid composition^35^. Their reduction during ZIKV infection therefore represents the previously unknown potential remodelling of a functionally relevant tripartite organelle contact. Further studies will be necessary to investigate the dynamics and mechanism of this reorganisation and their relationship to the changes in mitochondrial morphology observed in Fig. 5b.

Taken together, these data show that ZIKV-induced remodelling extends beyond individual compartments to a broad reorganisation of pairwise and higher-order relationships across the host cell organelle network. By recovering established infection-associated phenotypes, while identifying previously unrecognised changes in network organisation, PRISM demonstrates the potential of network-level organelle phenotyping to rapidly uncover host cell remodelling following pathogen infection, including in heterogeneous populations where only a subset of cells is infected.

## Discussion

In this study, we present PRISM, a method that combines stable five-organelle labelling with quantitative analysis of organelle phenotypes and network organisation. Encoding the complete reporter panel on a single PiggyBac-compatible construct allows the same compartments to be labelled consistently across cells and experiments, while retaining compatibility with additional molecular or functional probes for spectral imaging. We established stable PRISM lines in several cell types and validated the approach using cytoskeletal perturbations, recovering established organelle responses while identifying novel remodelling of pairwise and higher-order contacts. Furthermore, we demonstrate the combination of PRISM with additional fluorescent markers in the context of lysosomal damage and ZIKV infection, and show that it can reveal coordinated changes across the organelle network in individual cells.

Previous spectral imaging approaches have established the biological value of simultaneously visualising multiple organelles, and other multireporter constructs have demonstrated that several organelle labels can be expressed from a single vector^36, 37^. PRISM addresses the complementary challenge of reproducing a consistent labelling configuration across large cell populations and between experiments, and does so with a panel of markers that is optimised to robustly label organelles with minimal functional impact. Stable integration allows the complete reporter panel to be inherited as a single unit, while clonal selection can be used to establish uniform expression for reliable spectral separation. Once generated, the same labelled population of cells can be used across multiple perturbations without repeatedly introducing and optimising the organelle probes — a considerable challenge when working at scale. Importantly, the blue and far-red spectral regions remain available, allowing end users to add labelled proteins, lipids, dyes or genetically encoded fluorescent reporters without changing the underlying organelle panel.

The applications presented here illustrate how PRISM (Fig. 1) can support biological discovery. Following microtubule depolymerisation, we recovered a well-established Golgi fragmentation phenotype but also revealed lysosomal remodelling and changes in specific higher-order organelle relationships (Fig. 2,3). The addition of a farred biosensor allowed us to examine the molecular redistribution of PI(4)P following lysosomal damage alongside accompanying changes in lysosomal phenotypes, supporting the involvement of PI(4)P in damage-induced lysophagy (Fig. 4). In cells infected with ZIKV, we recovered known changes in ER, mitochondrial and peroxisomal characteristics while revealing a broader reduction in pairwise and higher-order contacts (Fig. 5), illustrating that PRISM can identify network-level responses that would not be apparent from analysis of one or two organelles in isolation.

As with any approach based on genetically encoded organelle reporters, expression of our exogenous organelle markers could influence the structures being measured. We selected targeting sequences and fluorescent proteins to minimise these effects, but reporter localisation and organelle morphology should be monitored when establishing new lines and during prolonged culture. Indeed, we note that peroxisomal dilation can occur in some PRISM lines at higher passage numbers. Stable expression may also be affected by promoter silencing in some cell types or during changes in cellular state, for example certain iPSC differentiations.

We implement our imaging and analysis in two dimensions to enable higher-throughput acquisition and imaging in both live and fixed cells. In relatively flat adherent cells, a single focal plane captures much of the dominant morphological and radial organisation of the labelled organelles, allowing larger cell populations and more experimental conditions to be compared. By excluding axial information, we could potentially miss vertical interactions between organelles situated in different focal planes. The regions we classify as contacts within the focal plane are defined by pixel overlap between diffraction-limited organelle masks. As such, they encompass both *bona fide* membrane contact sites as well as areas of close proximity between organelles. They should be viewed as an indication of where functional organelle interactions could be taking place, as well as candidates for subsequent validation using dedicated proximity reporters^38–40^.

The current analysis pipeline prioritises measurements that are entirely agnostic to broader morphology and consistent across organelles, so that they can be applied even in cases of drastic remodelling. Therefore, we do not implement more organelle-specific analyses, such as branching metrics. Where greater structural detail is required, the data from PRISM can be additionally analysed using dedicated packages such as ERnet or Mitometer^41, 42^.

Organelle network remodelling underpins changes in cellular behaviour and homeostasis in response to both extrinsic and intrinsic stimuli. PRISM makes it possible to reliably quantify these processes in stably labelled cell lines, allowing network-level responses to be compared directly across many experimental conditions. This creates opportunities, for example, to compare how different pathogens reshape the host organelle landscape, or to use organelle phenotypes as quantitative readouts for small-molecule or genetic screens. Given that PRISM spectra are compatible with additional blue and far-red functional reporters, proteins, lipid probes or responsive dyes, these organelle phenotypes can be linked directly to the biological processes under study. Specific changes identified by PRISM can then be further investigated mechanistically in targeted follow-up experiments. In summary, by labelling a consistent panel of organelles while accommodating additional experimental reporters, PRISM provides a reproducible framework for investigating organelle network remodelling across diverse biological contexts.

## Methods

### PRISM construct design and cloning

PRISM consists of five organelle markers and a puromycin resistance gene under three promoters, assembled into a PiggyBac transposon between inverted terminal repeats. Reporter sequences were cloned from existing plasmids or codon-optimised and sequenced *de novo*, as indicated in Tables 1 and 2 below. Promoters were cloned from plasmids gifted by Michael Ward; their corresponding expression cassettes are shown in Table 3. The IRES sequence and puromycin resistance gene were cloned from pLVX-EF1a-Neuromodulin-IRES-Puromycin *(Addgene #134666)*, which was a gift from David Andrews^43^. Further details are provided in Supplementary Text and Supplementary Fig. 1.

**Table 1.** Organelle targeting sequences.

| Organelle | Targeting sequence | Source |
| --- | --- | --- |
| Golgi | Giantin transmembrane region (3131–3258) | pmScarlet-Giantin-C1 (*Addgene #85048*), from Dorus Gadella<sup>44</sup> |
| lysosomes | TMEM192 full length | Synthesised by Integrated DNA Technologies |
| mitochondria | 2x COX8 mitochondrial localisation sequence | Synthesised by Integrated DNA Technologies |
| peroxisomes | SKL motif | Synthesised by Integrated DNA Technologies |
| ER | Calreticulin signal sequence; KDEL retention motif | Synthesised by Integrated DNA Technologies |

**Table 2.**
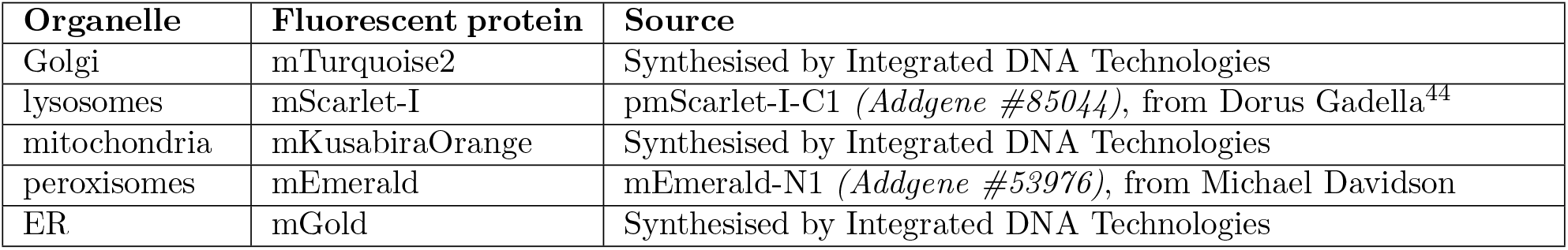
Fluorescent proteins.

| Organelle | Fluorescent protein | Source |
| --- | --- | --- |
| Golgi | mTurquoise2 | Synthesised by Integrated DNA Technologies |
| lysosomes | mScarlet-I | pmScarlet-I-C1 (*Addgene #85044*), from Dorus Gadella<sup>44</sup> |
| mitochondria | mKusabiraOrange | Synthesised by Integrated DNA Technologies |
| peroxisomes | mEmerald | mEmerald-N1 (*Addgene #53976*), from Michael Davidson |
| ER | mGold | Synthesised by Integrated DNA Technologies |

**Table 3.** Expression cassettes.

| Promoter | Marker 1 | Linking element | Marker 2 | polyA |
| --- | --- | --- | --- | --- |
| CAG | mitochondria | T2A | peroxisomes | bGH |
| EF1α | lysosomes | P2A | Golgi | bGH |
| hPGK | ER | IRES | puromycin resistance | rBG |

The full PRISM plasmid was cloned using sequential Gibson assembly. The mitochondrial and peroxisome reporters were assembled downstream of a CAG promoter, while the ER reporter and puromycin resistance gene were assembled under an hPGK promoter in another plasmid. The coding sequences from each of these were amplified by PCR using Gibson assembly primers (Table 4) and inserted into the PiggyBac backbone, which was linearised with BsrGI-HF and KpnI-HF *(New England Biolabs, R3575 and R3142, respectively)*. The resulting plasmid was then linearised with BmtI and SwaI *(New England Biolabs, R0658 and R0604, respectively)*, while the final segment was amplified using Gibson assembly primers (Table 4) from another plasmid containing the assembled Golgi and lysosome reporters to make the final PRISM construct. All assembly reactions were carried out using NEB HiFi Master Mix *(New England Biolabs, E2621S)*. The resulting plasmids were transformed in NEB Stable competent *E. coli (New England Biolabs, C3040I)* and purified using the Qiagen Plasmid Plus Maxi kit *(Qiagen, 12963)*. The final plasmid was verified by Sanger sequencing.

**Table 4.**
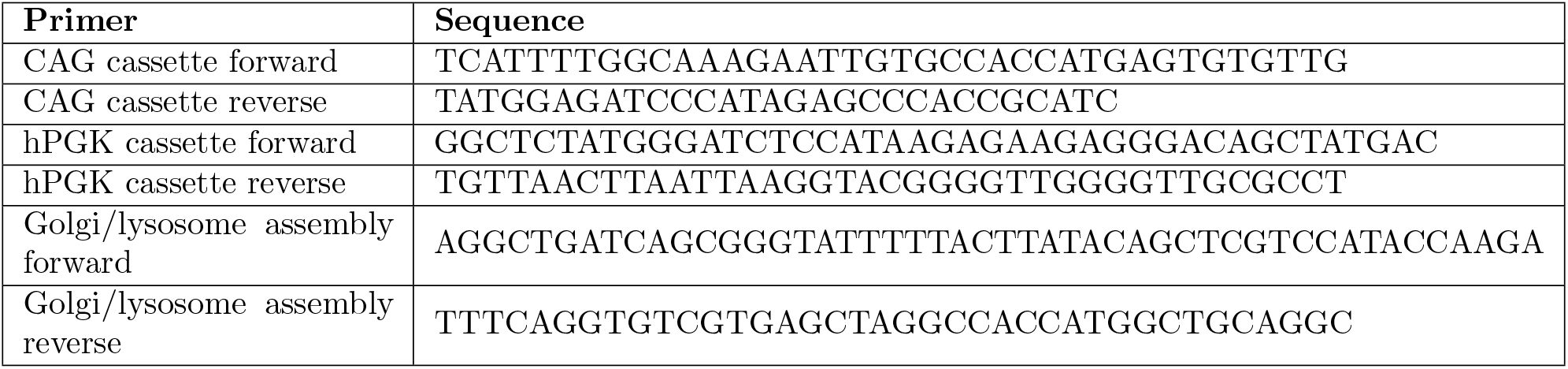
Gibson assembly primers.

#### Other plasmids

Single-colour control plasmids were used to assess the individual organelle markers and to collect reference spectra for image unmixing. These were cloned from the same sources listed above into a standard plasmid containing an EF1α promoter. A pCC1BAC vector containing the open reading frame of an Asian-American lineage ZIKV with an mCherry tag fused to the capsid protein was kindly provided by Andres Merits^45^. The mCherry was replaced with the HaloTag from HaloTag-N1, a gift from Jennifer Lippincott-Schwartz. A plasmid encoding the Super PiggyBac transposase enzyme under the control of EF1α was kindly provided by Michael Ward.

### Cell culture

HeLa, U2OS, and COS-7 cell lines *(ATCC; CCL-2, HTB96, and CRL1651, respectively)* were maintained in DMEM *(Sigma, D6546)* with 10% FBS *(Gibco, A5256701)*, 100U/mL PenStrep *(Merck, P4333)*, and 1x GlutaMAX *(Gibco, 35050061)*. Vero cells *(ATCC, CCL-81)* were grown in DMEM supplemented with supplemented with 100U/mL penicillin, 100 µg/mL streptomycin, and 2 mM L-glutamine. All cells were grown in a humidified incubator at 37°C with 5% supplemental CO_2_. Cell lines were tested monthly for mycoplasma contamination by PCR and were consistently negative.

#### PiggyBac integration

Stable PRISM-expressing HeLa, U2OS, and COS-7 lines were generated by co-transfecting the PRISM and Super PiggyBac transposase plasmids at a mass ratio of 2:1 using FuGENE HD *(Promega, E2311)*. For each transfection, 1 µg PRISM and 0.5 µg transposase DNA were added to cells in a single well of a 6-well plate. 48 hours after transfection, cells were selected in medium containing 2.5 µg/mL puromycin (10 µg/mL for U2OS cells) for one week, after which they were expanded in growth media without puromycin. The HeLa line used in this paper also contains an integrated Zim3-dCas9 with a nuclear mTagBFP2 reporter to facilitate subsequent CRISPRi experiments. mTagBFP2 was not included as a component during spectral unmixing; residual signal in the mTurquoise2 (Golgi) channel was excluded during segmentation and was not included in further quantitative analysis.

#### Clonal line selection

The initial polyclonal PRISM HeLa line was sorted using a Thermo Fisher Bigfoot Spectral Cell Sorter. Single cells were sorted into a 96-well plate containing complete growth media supplemented with an additional 10% HI-FBS *(Gibco, A5256801)* and 1% CloneR2 *(STEMCELL Technologies, 100-0691)*. Once colonies had grown, they were transferred to a 96-well glassbottom imaging dish *(Cellvis, P96-1*.*5H-N)* for spectral imaging. The clones were assessed for reporter expression, spectral unmixing, growth, and organelle morphology. A single clone was selected and used for all subsequent experiments.

### Drug perturbations

For microtubule disruption experiments, HeLa cells were placed in cold media with the addition of either 5 µM nocodazole *(abcam, ab120630)*, or 0.1% (v/v) dimethyl sulfoxide, and incubated on ice for 2 minutes to induce depolymerisation of existing microtubules. Cells were then transferred to 37°C with 5% supplemental CO_2_ to recover for 1 hour before imaging under the same conditions. Nocodazole or DMSO remained present in the media during acquisition, which was completed within 1 hour.

For actin disruption, cells were placed in 37°C media containing either 1 µM cytochalasin D *(Cayman Chemical, 11330)* or 0.025% (v/v) DMSO. These were incubated at 37°C with 5% CO_2_ for 1 hour before imaging under the same conditions. Cytochalasin D or DMSO remained present in the media during acquisition, which was completed within 1 hour.

To induce lysosomal damage, cells were treated for 1 hour with media containing 0.5 mM LLOMe *(Sigma-Aldrich, L7393)* or 0.1% (v/v) DMSO, and fixed immediately afterwards as described below.

### ZIKV production and infection

The Halo-tagged virus construct was linearised with AgeI *(New England Biolabs, R3552)* and transcribed *in vitro* using the mMESSAGE mMACHINE SP6 Transcription Kit *(Thermo Fisher, AM1340)*. Capped RNA was electroporated into Vero cells as previously described^45^. Passage 0 (P0) supernatant was collected after a significant cytopathic effect was observed, typically at approximately six days post-electroporation. This was then used to infect more Vero cells to produce the P1 stock. P1 supernatant was collected three days after infection and concentrated by ultracentrifugation through a layer of 25% glycerol-TNE buffer (100 mM Tris-HCl pH 8, 150 mM NaCl and 1 mM EDTA) at 110,000 x g for 3 hours at 4°. The ZIKV pellet was resuspended in TNE buffer, aliquoted, and stored at −80°.

ZIKV infectivity was assessed with tissue culture infectious dose (TCID_50_) assays as described in^46^. Viral stocks were subjected to 10x serial dilution and used to infect cells in a 96-well plate. Cells were incubated at 37°C for 5 days before being fixed with 4% formaldehyde and 0.9% NaCl for 30 minutes. Cells were stained with 0.1% toluidine blue, and wells exhibiting a cytopathic effect were scored as positive. TCID_50_ values were calculated using the Reed-Muench method. Plaque-forming units per millilitre (PFU/mL) were estimated from the TCID_50_ values using a conversion factor of 0.693^47^.

30 hours prior to infection, PRISM-labelled HeLa cells were seeded onto 13 mm glass coverslips in a 24-well plate at a density of approximately 7500 cells per well and grown in serum-free DMEM supplemented with 100 U/mL penicillin, 100 µg/mL streptomycin, and 2 mM L-glutamine. After 24 hours, this was replaced with media supplemented with 5% FBS. Six hours later, cells were infected with Halo-tagged ZIKV in serum-free media at an MOI of 0.04. The virus was left on the cells for 1 hour at 37°C, then replaced with DMEM supplemented with 2% FBS and 20 mM HEPES at pH 7.4. For the uninfected condition, cells underwent identical media changes, except with serum-free DMEM in place of the virus.

### Fixation/staining protocols

#### PI4P staining

The Alexa Fluor 647-labelled PI(4)P biosensor was a kind gift from Hannes Maib. Cell fixation and staining were carried out as described in Maib et al. 2024^48^. DMSO control or LLOMe-treated cells were fixed for 20 minutes at room temperature with 4% formaldehyde and 0.2% glutaraldehyde in PBS before rinsing with PBS and quenching for 20 minutes in 50 mM NH_4_Cl. They were then placed on ice and rinsed with cold PIPES buffer containing 10 mM piperazine-N,N’bis[2-ethanesulfonic acid], 10 mM NaCl, and 1 mM MgCl_2_. The PI(4)P biosensor was then added at 500 nM in PIPES buffer with 5% bovine serum albumin (BSA) and 0.5% saponin, and incubated for 1 hour. Cells were then washed, post-fixed with 2% formaldehyde in PBS, and finally quenched again with NH_4_Cl.

#### ZIKV fixation/Halo labelling

35 hours after infection, the JFX646 HaloTag ligand (a gift from Luke Lavis) was applied to cells at a dilution of 1:1000 in OptiMEM *(Gibco, 31985062)* for 1 hour. Subsequently, cells were washed twice with PBS before fixing with 4% PFA and 0.2% glutaraldehyde for 15 minutes at room temperature before quenching in 15 mM glycine for 20 minutes, and washed with PIPES buffer as described above. Cells were then stained with DAPI (1:10k) for ten minutes before mounting on glass slides with ProLong Gold antifade medium *(Thermo Fisher, P10144)*.

### Imaging

For drug treatment experiments, HeLa cells were plated the day before imaging to a final confluence of 10–20% on glass coated with Matrigel *(Corning, 356231)*. Depending on the experiment, either 25 mm #1.5 coverslips *(Scientific Laboratory Supplies, MIC3350)* or 8-well imaging dishes *(ibidi, IB-80807)* were used. For ZIKV experiments, cells were plated on 13 mm glass coverslips as described above.

All imaging in this paper was carried out on a Zeiss LSM980 laser scanning confocal, running on ZEN Blue 3.12. Images were acquired using a 63x/1.40 Plan-Apochromat DIC M27 oil immersion objective and a 32-channel GaAsP-PMT detector array (411–639 nm range), with PRISM-labelled cells excited by 405, 514, 561, and 639 nm lasers.

Single optical sections were acquired with 52 µm pinhole, 1.20x sampling (optimised for LSM Plus), and twofold line averaging. Images were acquired at 16-bit depth with a frame size of 2290×2290 pixels and a pixel dwell time of 1.83 × 10^−6^ seconds. The final pixel size after LSM Plus processing was 0.059 µm. Laser powers and detector settings were held constant within each experiment.

The example images in this paper have had their brightness and contrast adjusted for clarity; quantitative analyses were performed on unadjusted images. Lookup tables used for composite images are detailed in Supplementary Text.

### Image analysis

#### Unmixing and deconvolution

Reference spectra were collected using the same microscope configuration from stable HeLa lines expressing the individual organelle markers described above. Reference spectra for DAPI, JFX646, and the Alexa Fluor 647-conjugated PI(4)P biosensor were acquired from labelled parental HeLa cells. For drug treatment experiments, these spectra were used to perform linear unmixing in ZEN, with residuals. These residuals were included in the maximum intensity projection used for cell boundary segmentation, but otherwise excluded from further analyses. For ZIKV experiments, the spectra were used to set up an Online Fingerprint, and unmixing was performed during acquisition. Unmixed images were deconvolved using LSM Plus Processing in ZEN with default settings.

#### Segmentation

Deconvolved CZI images were converted into single-channel TIF files for organelle segmentation. Organelle masks were generated by supervised pixel classification in ilastik 1.4.1. For each experiment, a separate classifier was trained for each organelle on 2–3 images from each experimental condition. Performance was assessed on a similar number of test images, and any poorly segmented images were added to the training set for additional training. The final classifiers were applied to all images from that experiment to produce binary segmentation masks. For nucleus segmentation from DAPI staining in the LLOMe experiment, an additional ilastik object classification step was performed to apply smoothing and size thresholding. Cells with no nucleus segmented or with obvious errors in organelle segmentation were excluded from further analysis.

For the nocodazole and cytochalasin D experiments, cell and nucleus masks were traced manually in ImageJ 1.54p using a maximum intensity stack of all unmixed channels. For the LLOMe experiment, an ilastik pixel classifier was trained as above to identify cells from the maximum intensity projections, and the binary output of this classifier was then manually processed in ImageJ and GIMP 3.08+ to separate abutting cells, assign unique pixel values to each cell in an image, and fill in small holes in the segmentation. For ZIKV experiments, cells and nuclei were segmented using a custom CellProfiler 4.2.8 pipeline. Cells intersecting an image boundary were excluded from the segmentation.

#### Custom software

To facilitate the analyses described in this paper, we wrote two custom ImageJ macros and an RMarkdown notebook, which are available at github.com/JNA-Lab/Organelle analysis. Their function is described in more detail in the Supplementary Text, and a user guide is available on the GitHub page. These form a pipeline to process segmented cell and organelle masks into 507 quantitative metrics of organelle size, shape, position, and contacts within the cell. Contacts are calculated by pixel overlap between organelle masks; as these masks reflect the diffraction limit of the original data, we do not distinguish closely apposed membranes from *bona fide* membrane contact sites, but refer to them as ‘contacts’. Spatial metrics calculated include mean distribution radius (MDR) and coefficient of asymmetry (CoA), which we have adapted from Zheng et al. 2022^11^. Details of our implementation are in the Supplementary Text. CoA in this paper was calculated with a wedge angle of 5°. Optionally, the pipeline can also quantify fluorescence intensity from any given channel within each cell, organelle, and contact mask and combine this with the basic organelle metrics to allow both to be assessed in the same cells. The final output is a data frame in R, which can be used directly for further analysis and visualisation or exported as a CSV.

### Data normalisation and analysis

All reported organelle areas are normalised to the corresponding cell area. Organelle contact areas are normalised to the cumulative area of the involved organelles; in other words, if the two (or more) organelles completely overlap, the normalised contact area will be 1, regardless of their relation to the total cell area.

Downstream analysis and data visualisation was carried out using R 4.5.3^49^. The ‘ggplot2’ package version 4.0.2^50^ was used for plotting, with continuous colour maps from Fabio Crameri^51^. Statistics were carried out with the base R ‘stats’ package and ‘ggpubr’ version 0.6.3^52^.

## Statistics

For comparisons across multiple metrics (e.g. Fig. 2c, d), individual cell measurements were treated as observations. Two-sided, unpaired Wilcoxon rank-sum tests were used to compare metric values between control and experimental conditions, and *p*-values were adjusted for multiple comparisons with the Benjamini-Yekutieli method. Additionally, Z-scores for each metric were calculated relative to the mean and standard deviation of the control condition, and mean zscores were plotted for visualisation purposes.

Individual metrics were selected for further analysis based on the above global statistics or biological relevance, and were analysed treating the mean of each experimental replicate as an observation. For drug treatment experiments with two experimental conditions, two-sided Welch’s t-tests were used to compare the means of each condition. For the ZIKV experiment, ordinary one-way ANOVA was performed, and where it returned a *p*-value of *≤* 0.05 it was followed up with post-hoc twoway t-tests for all possible pairs, with Bonferroni correction for multiple comparisons. No statistical method was used to predetermine sample size, and experimenters were not blinded to condition during the analysis.

## Acknowledgments

The authors would like to acknowledge the CIMR Core Facilities staff for their contributions to this work. In particular, we would like to thank Matthew Gratian and Mark Bowen for microscopy assistance, and Reiner Schulte and Gabriella Grondys-Kotarba for cell sorting. We would also like to acknowledge the University of Cambridge Statistics Clinic for their advice on the analyses in this paper.

## Declarations

### Data availability

All data presented in this paper are available by request.

### Materials availability

The PRISM plasmid and cell lines produced are available by request; please contact Jonathon Nixon-Abell.

### Code availability

The organelle analysis code produced in the course of this work is available on GitHub at github.com/JNA-Lab/Organelle analysis. Code used to generate the figures in this paper is available by request.

### Generative AI use

During the preparation of this manuscript, JNA used Claude Code for copy editing. The authors have reviewed and edited the material and take full responsibility for the final paper.

### Author Contributions

**FML:** Conceptualization, Data curation, Formal analysis, Investigation, Methodology, Software, Visualization, Writing - original draft, Writing - review & editing **ZB:** Data curation, Formal analysis, Investigation, Software **ST:** Software **TA:** Software, Visualization **HC:** Investigation **NA-D:** Investigation **SW:** Investigation **HM:** Resources, Methodology **NI:** Conceptualization, Funding acquisition, Project administration, Supervision **JNA:** Conceptualization, Funding acquisition, Investigation, Project administration, Resources, Supervision, Writing - review & editing

