## Supplementary Materials for "PRISM: A Plasmid-based Reporter for Intracellular Spectral Microscopy"

### Supplementary Figures

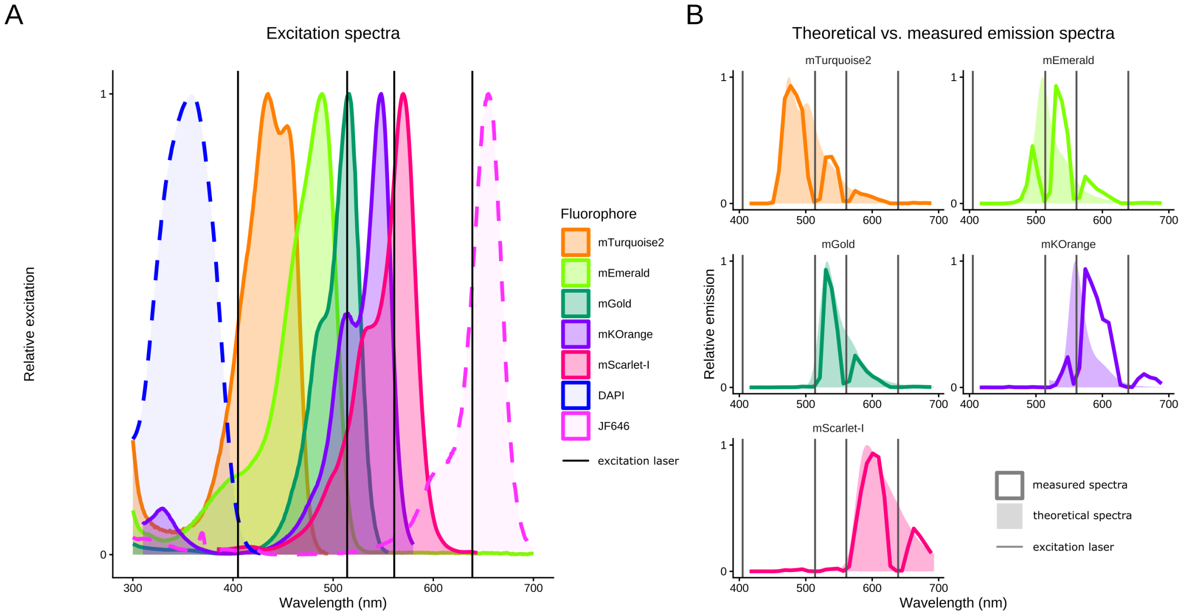


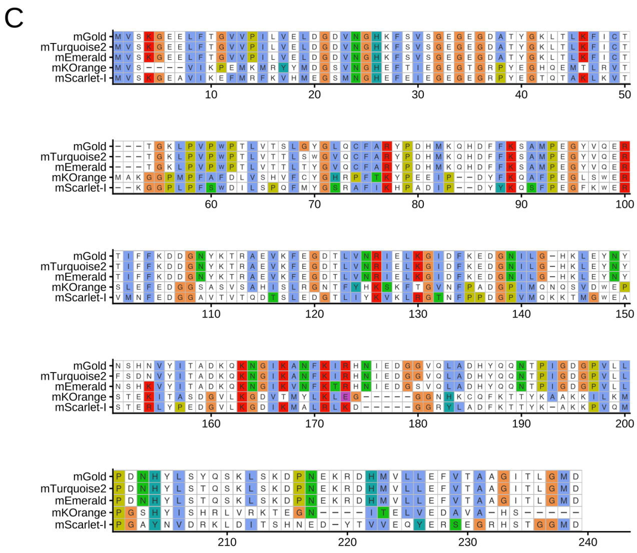


D

| Protein | OSER score | Source |
|---|---|---|
| mEmerald | 96.6 ± 1.1 | Cranfill et al. (2016) |
| mGold | 79.3 ± 6.7 | Lee et al. (2025) |
| mKOrange | 46.9 ± 6.1 | Cranfill et al. (2016) |
| mScarlet-I | 76 | Bindels et al. (2016) |
| mTurquoise2 | 93.8 ± 1.0 | Cranfill et al. (2016) |

**Supplementary Figure 1.** **(a)** Excitation spectra of the fluorescent proteins included in PRISM alongside example blue (DAPI) and far-red (JF646 Halo ligand) reporters as used in Fig. 5. Vertical lines indicate the excitation lasers used for imaging. Colours match the LUTs used throughout the manuscript. **(b)** Theoretical vs. measured emission spectra for the fluorescent proteins used in PRISM, as measured in live cells. Vertical lines indicate the excitation lasers used for imaging; these wavelengths are blocked from the collected fluorescence data by metal pins, hence the 'notches' in the observed spectra. This accounts for the apparent red shift in the mEmerald and mKOrange spectra, where the 'peak' fluorescence wavelength is not collected because it coincides with an excitation laser. **(c)** ClustalW amino acid alignment of the fluorescent proteins included in PRISM. **(d)** Table of OSER oligomerisation scores for fluorescent proteins included in PRISM[1–3].

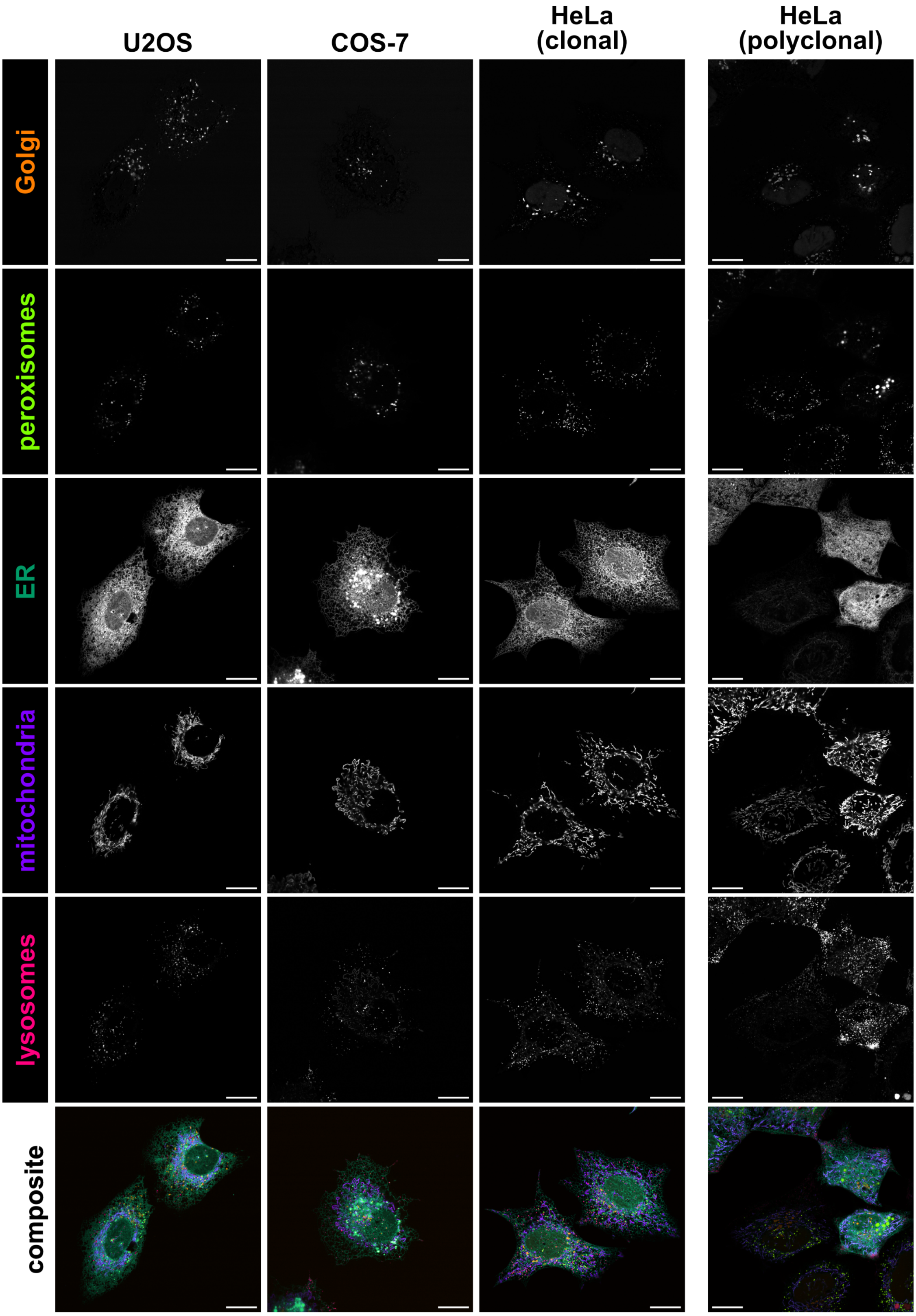


**Supplementary Figure 2.** Individual channels of representative U2OS, COS-7 and HeLa PRISM cells shown in Figure 1. Righthand panels show a polyclonal HeLa line — note the variation in brightness between cells. Both HeLa lines contain a nuclear-localised mTagBFP2 as a marker for integrated Zim3-dCas9. This signal is not unmixed from the other organelle channels and thus is slightly visible in the Golgi channel prior to its removal during segmentation. Images representative of cells from at least 3 independent experiments. Scale bars, 20 µm.

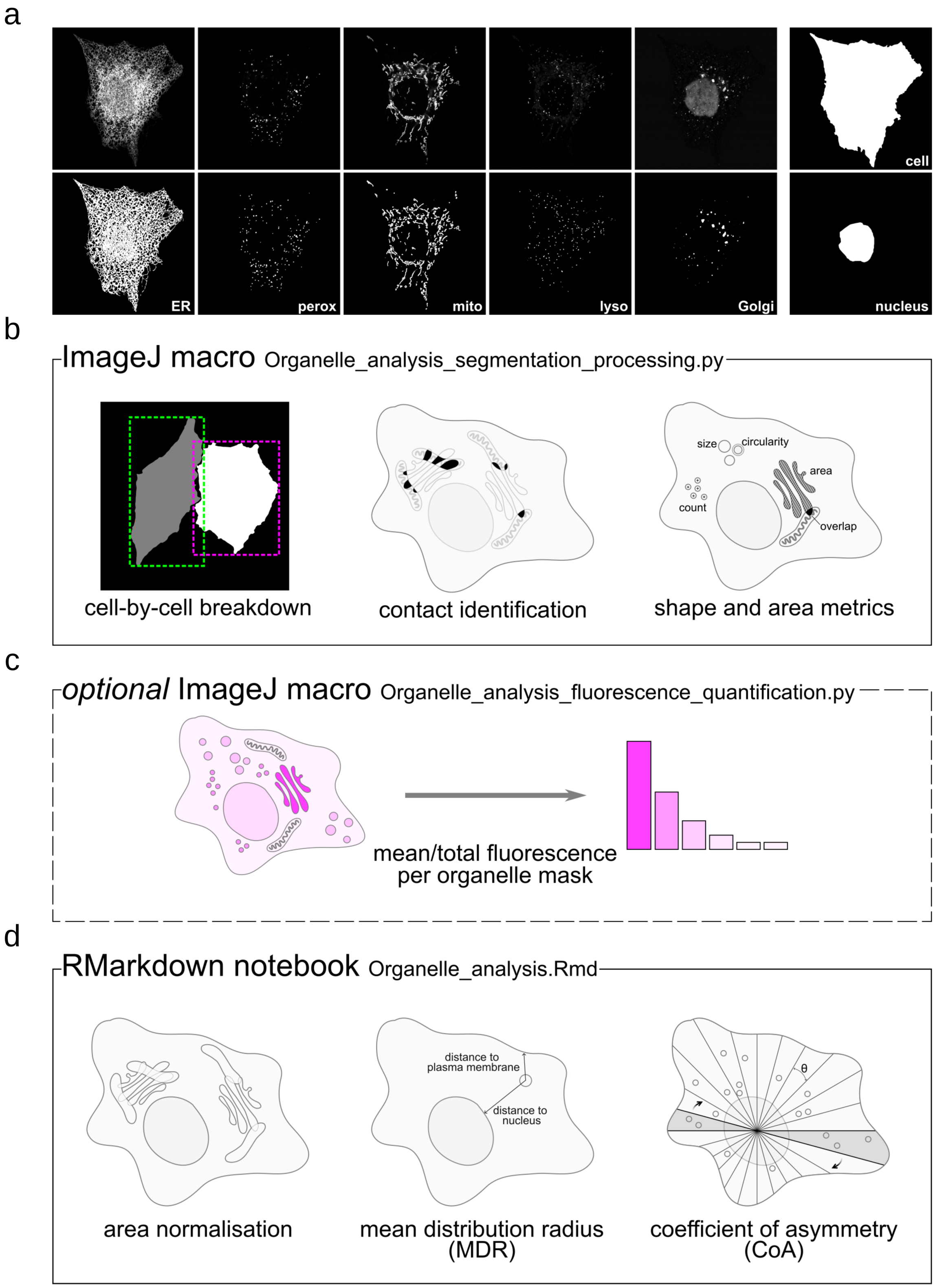


**Supplementary Figure 3.** **(a)** Individual channels of a representative HeLa PRISM cell with its corresponding segmentation masks generated from ilastik. The parental HeLa line contains a nuclear-localised mTagBFP2 as a marker for integrated Zim3-dCas9. This signal is not unmixed from the other organelle channels and thus is slightly visible in the Golgi channel prior to its removal during segmentation. Manually generated cell and nuclei masks are displayed alongside. Image shown representative of cells from 3 independent experiments. **(b-d)** Step-by-step overview of the standard PRISM image analysis pipeline (see Supplementary Text for details).

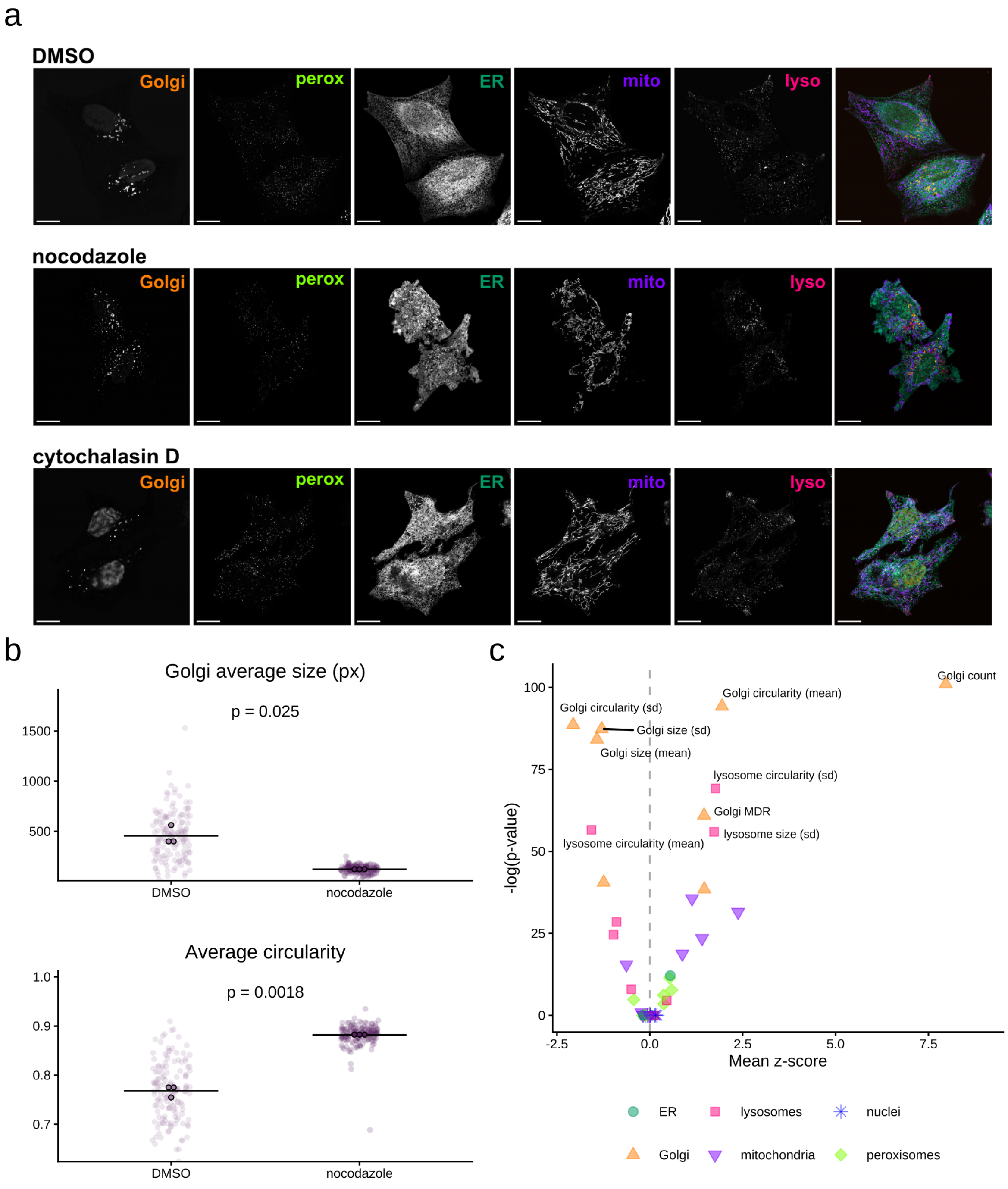


**Supplementary Figure 4.** **(a)** Individual channels of representative PRISM HeLa cells shown in Fig. 2a treated with DMSO, 5 µM nocodazole or 1 µM cytochalasin D for 1 hour. The parental HeLa line contains a nuclear-localised mTagBFP2 as a marker for integrated Zim3-dCas9. This signal is not unmixed from the other organelle channels and thus is slightly visible in the Golgi channel prior to its removal during segmentation. Images representative of cells from 3 independent experiments. Scale bars, 20 µm. **(b)** Plots of Golgi size and circularity in nocodazole-treated vs DMSO control cells. *P*-values from unpaired two-sided Welch's t-test. **(c)** Volcano plot of the nocodazole data presented in Fig. 2d, with *p*-value on the y-axis and mean z-score on the x-axis. Note that the most significantly changed metrics, after Golgi-related measurements, are related to lysosome size and shape. *P*-values from a Wilcoxon rank-sum test with Benjamini-Yekutieli correction for multiple comparisons.

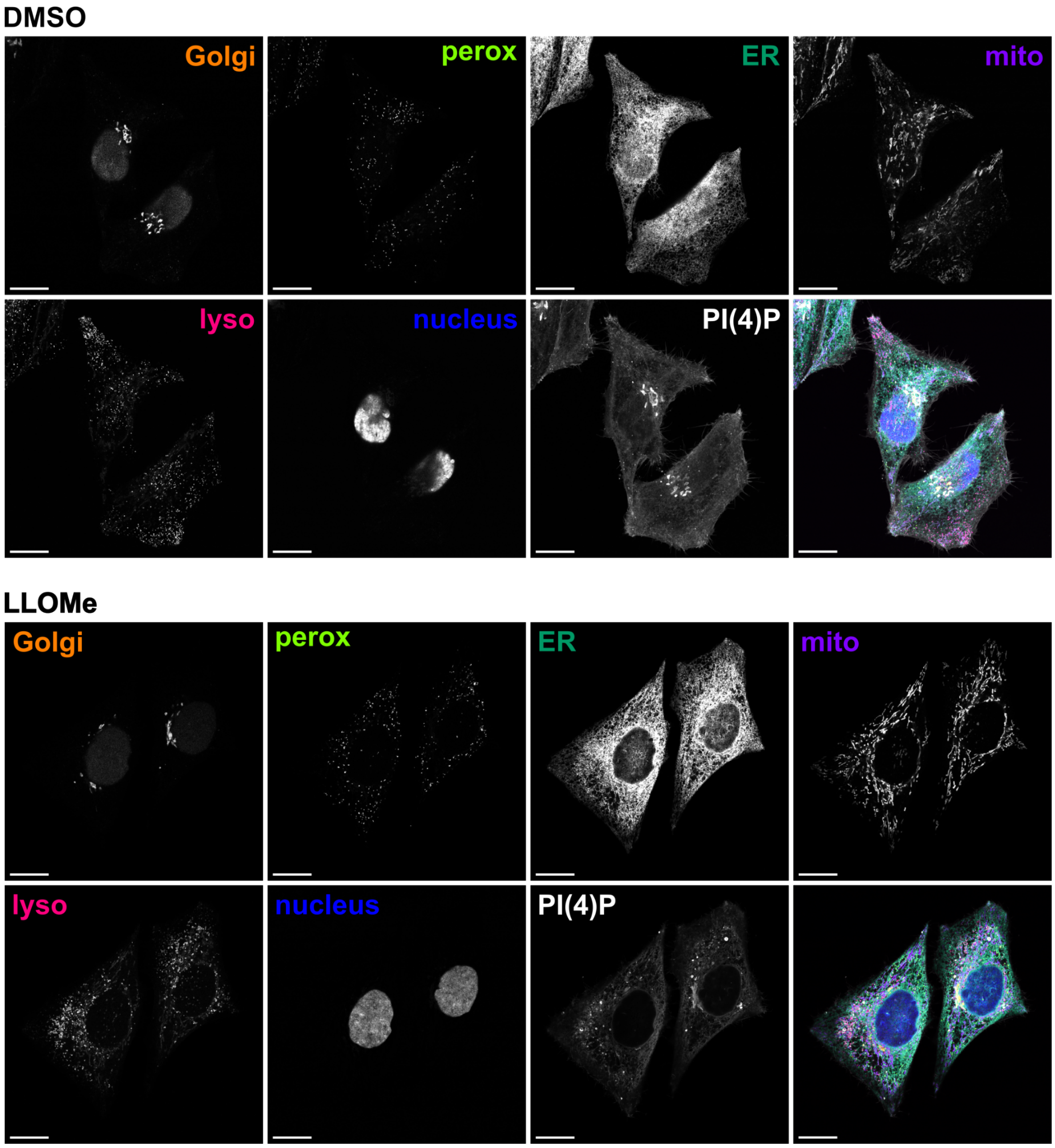


**Supplementary Figure 5.** Individual channels of representative PRISM HeLa cells shown in Fig. 4a treated with DMSO or 0.5 mM LLOMe for 1 hour, fixed, and co-stained with DAPI and an AF647-conjugated PI(4)P biosensor. The parental HeLa line contains a nuclear-localised mTagBFP2 as a marker for integrated Zim3-dCas9. This signal is not unmixed from the other organelle channels and thus is slightly visible in the Golgi channel prior to its removal during segmentation. Images shown are representative of cells from 3 independent experiments. Scale bars, 20 µm.

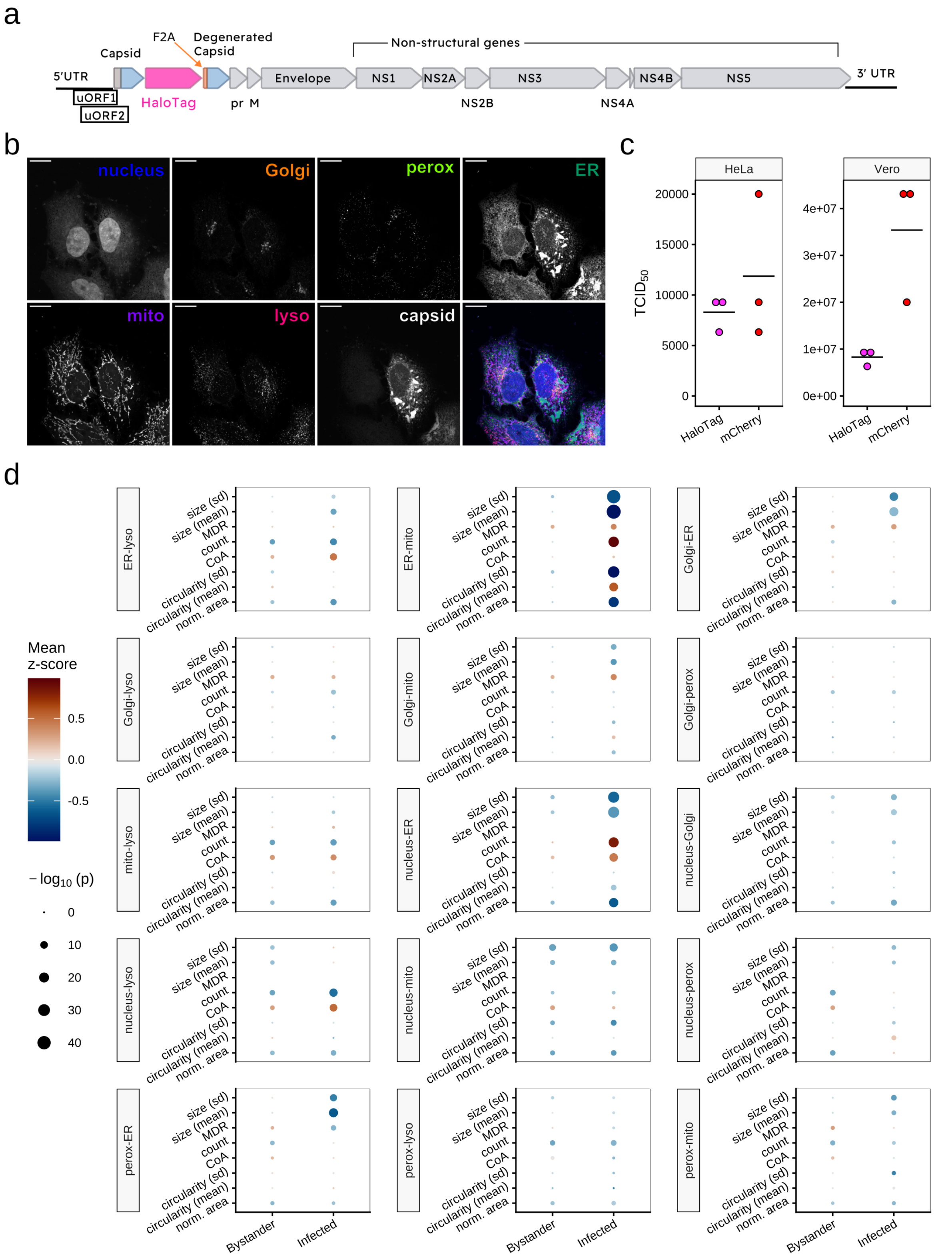


**Supplementary Figure 6.** **(a)** A linear map of the edited Zika virus genome including the Halo-tagged capsid. NS - Non-structural protein. **(b)** Individual channels of representative PRISM HeLa cells on a ZIKV-infected coverslip from Fig. 5a. The righthand cell is infected with the virus, while the lefthand cell is an uninfected 'bystander'. Images shown are representative of cells from 4 independent experiments. Scale bars, 20 µm. **(c)**

Tissue culture infectious dose ($TCID_{50}$) comparison data for HaloTag- and mCherry-tagged[4] ZIKV in PRISM HeLa cells and unlabelled Vero cells. n = 3 independent experiments. **(d)** Dot plots showing changes in organelle contact metrics between bystander and ZIKV-infected cells vs uninfected control cells. Point size indicates the adjusted *p*-value from a Wilcoxon rank-sum test with Benjamini-Yekutieli correction for multiple comparisons, colour indicates the mean z-score for each metric relative to the uninfected control. n = 263 (bystander), 289 (infected), 630 (uninfected) cells from 4 independent experiments.

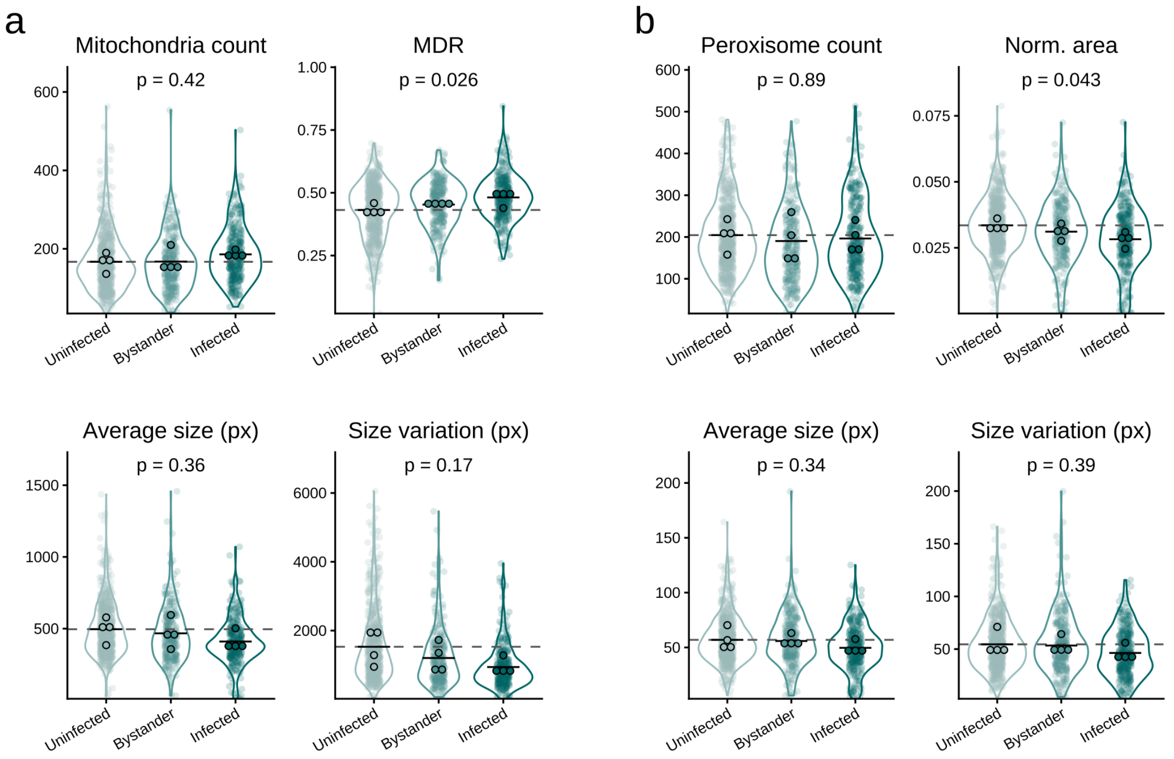


**Supplementary Figure 7.** **(a)** Plots of mitochondrial count, MDR, average size, and size variation (standard deviation) across uninfected (control), bystander, and ZIKV-infected cells. **(b)** Plots of peroxisome count, area, average size, and size variation (standard deviation) across uninfected (control), bystander, and ZIKV-infected cells. Mean = solid black line, comparison to uninfected mean = dashed gray line. *P*-values reported are from ANOVA. Where $p < 0.05$ pairwise Welch's t-tests (two-sided, unpaired) with Bonferroni correction for multiple comparisons were performed on independent replicate means, n = 4, but all results were non-significant.

These statistics are more conservative than those performed on individual cell values in Fig. 5b, which returned significant *p*-values for mitochondrial count (1.75e-4), MDR (1.39e-7), size (1.18e-8), and size variation (1.92e-20), peroxisome area (7.23e-8), size (4.28e-4), and size variation (2.14e-4) in infected cells, as well as mitochondria size variation (2.93e-4) and peroxisome count (3.58e-3) in bystander cells. Nonetheless, the trends we see match those reported in the literature, despite studying an earlier timepoint post-infection (36 hours vs 48 hours[5–7]).

### Supplementary Text

#### PRISM plasmid design

Our goal in developing PRISM has been to make spectral organelle imaging accessible and reproducible. Accordingly, we have optimised the plasmid to robustly label our selected organelles — Golgi, peroxisomes, ER, mitochondria, and lysosomes — without interfering with their normal function, morphology, or interactions. This drove our selection of organelle targeting sequences and associated fluorescent proteins.

Wherever possible, we used minimal targeting motifs for each organelle[8–12]. The only full-length protein used is TMEM192, which has been shown to label a broad population of lysosomes with more specificity than other commonly used reporters such as LAMP1[13,14].

The fluorescent proteins were selected based on several criteria. First, they required sufficiently distinct emission peaks for effective unmixing when imaged with a spectral detector array. The proteins we selected have an average peak separation of just under 30 nm, which should be sufficient to discriminate on a standard detector array. Second, they all needed to be monomeric, to avoid any dimerisation-induced interactions or morphological changes to the labelled organelles. Of these, mKOrange has the lowest OSER score, and so was selected for a marker that localises to the organelle lumen (mitochondria) rather than the membrane, to avoid any dimerisation-induced artefacts. Finally, they needed to be spectrally compatible with other fluorescent markers to enable imaging in combination with additional proteins, lipids or biosensors within the cell. We decided to leave the blue (<450 nm) and far red (>630 nm) spectral range free for compatibility with commonly used reporters, with capacity for multiplexing with near infrared reporters in the future. The selected panel of fluorescent proteins meet these criteria, and we have found them to be sufficiently bright and stable in combination with our organelle-targeting sequences.

Due to the inclusion of flanking PiggyBac ITR sites, co-transfecting PRISM with a plasmid containing the PiggyBac transposase will drive insertion into the cell genome, with the average number of insertions per cell tunable by varying the relative amounts of the two plasmids. After 48 hours, cells that have successfully integrated the PRISM construct can be selected with puromycin and expanded. Due to the nature of the PiggyBac transposon-transposase integration, these lines can be somewhat heterogeneous, as each cell may have a different number of genomic integration sites. While this heterogeneity is not a problem for most use cases, we find that it can decrease the accuracy of the image unmixing process. To address this in our HeLa line, we selected a clonal line through FACS, as described in Methods and illustrated in Fig. 1. We gated to select cells with good expression of all markers, but avoiding excessively bright mEmerald and mKOrange (both expressed under the CAG promoter) to avoid a peroxisomal dilation artefact associated with extended overexpression of SKL-tagged proteins. This process resulted in a more visibly uniform population and a marked increase in unmixing performance.

#### Image analysis

To facilitate analysis of 2D PRISM imaging data, we wrote two ImageJ macros and an RMarkdown notebook. They operate on *segmented* data, rather than directly on fluorescence images.

We designed these tools to be as flexible and customisable as possible while keeping them simple to use with minimal technical knowledge required. They do not require installation; they can simply be dragged and dropped into Fiji or RStudio and run directly. They will work on any subset of the organelle channels provided by the PRISM system (or labelled by alternative means), with the option to add an additional segmented organelle channel for other structures of interest within the cell (e.g. additional organelles, pathogens, etc.). The only strict requirement is the provision of cell masks, which are used for many of the downstream processing steps, plus at least one organelle channel, although we highly recommend the inclusion of nucleus masks. In the absence of a nucleus channel, the MDR and CoA metrics described below will be calculated based on the cell centroid, which may be less accurate.

These tools also provide the means to extract metadata from image file names. The file names are split at each underscore, and each piece of the name can then be assigned to a particular variable — for example an image entitled "HeLa_replicate3_control_image2" could be assigned to variables for cell type, experimental replicate, condition, and image number, which would all be extracted automatically and available for downstream analysis.

##### ImageJ macros

The function of the main ImageJ macro is to break the segmented images down into individual cells, calculate the pixel overlap between organelles, and calculate shape and size measurements for each organelle or contact. It also outputs a number of additional files that can be used for optional extra steps (e.g. fluorescence quantification, described below), or alternative analysis tools.

This process begins by matching up the segmentations corresponding to each individual image. This uses the file names, plus the unique suffixes for each channel specified in a user input dialog.

**Overlap calculation** The macro computes all possible combinations of the organelles present (both pairwise and 'higher order', e.g. three- and four-way contacts) and creates pixel overlap masks for all of them. These are then stacked with the 'base' organelle channels and treated in the same way in further processing steps, so all the same metrics are available for contacts as for organelles.

As this pipeline is intended to operate on (segmented) confocal imaging data, with resolution constrained by the diffraction limit of the microscope used, we cannot say that these contacts directly correspond to functional membrane contact sites in the cell. However, they set an upper limit on the possible area of membrane contact sites within the imaged region of the cell, and indicate where these interactions could be located in relation to the rest of the organelle network.

**Cell separation** With this stack of organelle and contact channels created, it is then cropped using the segmented cell masks to produce a separate stack for each cell in the image. This stack is used for subsequent analysis steps to calculate organelle metrics for each cell.

**Measurements - individual, pooled, and summary** First, particle analysis is performed on each organelle channel within the cell. This produces an ROI for each separated organelle, along with associated measurements such as size and circularity. This step is carried out on *binary* masks of the organelle, so overlapping particles are treated as a single organelle. Then, the organelle ROIs within each channel are combined, to give the total area of each organelle within the cell as well as ROIs for later processing steps. Finally, all organelle and all contact ROIs are combined to give the summary metrics for each cell — the total area of the cell occupied by all organelles or organelle contacts.

**Output files** The macro outputs two files for each original image, plus an additional seven files for each cell within the image. These files generally do not need to be handled directly, as the next steps in the processing pipeline will simply accept the path to the analysis folder and handle retrieving data from the correct files, but they are available for alternative analyses if desired.

For each image, two ROI ZIP files are produced, one with cell mask ROIs (`..._ALL_CELL_ROIs.zip`) and another (`..._ALL_ORGANELLE_ROIs.zip`) that additionally contains organelle ROIs (one per organelle type) for each cell within the image. These are intended for additional analyses on the original images, including fluorescence quantification (for which we have provided an additional macro, detailed below).

For each cell, a TIF stack is produced from the cropped masks. This file is used for calculating additional organelle distribution metrics in R, but also makes it possible to quickly check any outliers that emerge later in the analysis. This is accompanied by two ROI files, one with each individual organelles annotated (`..._ROIs.zip`), and another (`..._POOLED_ROIs.zip`) with the organelles pooled together by type (so that e.g. all peroxisomes will be a single ROI). These are not used directly in our metrics, but are available to be used for other analyses — for example organelle-specific morphology characterisation of the ER or mitochondria. Finally, there are four CSV files produced for each cell — one for each of the individual, pooled, and summary measurements calculated above (`..._results.csv`, `...POOLED_results.csv`, and `..._summary_results.csv`, respectively), and one small file containing the slice names of the TIF stack

(`..._slice_labels.csv`), which is used to distinguish organelle and contact channels later in the analysis process. For users concerned about data storage space, the cell TIF stacks are by far the largest files output from this macro. They are required for the following RMarkdown file to run, but can be deleted afterwards if necessary.

**Optional - fluorescence quantification macro** A separate macro is provided to quantify the fluorescence intensity of defined channels within each organelle mask. This uses the above mentioned `..._ALL_ORGANELLE_ROIs.zip` to map the identified organelles onto the original fluorescence images to calculate the mean and total fluorescence within each organelle on a per-cell basis, for user-defined channels. This data is output as one CSV file per image and can be directly read in by the RMarkdown notebook at the next step to combine this data with the other metrics.

##### RMarkdown notebook

The RMarkdown notebook consolidates data from the multiple files output by the Images macro(s), computes additional spatial metrics, normalises data, and filters the final metric table to remove redundant or unnecessary measurements. The final output is a table (as a data frame in R) that can be used directly for downstream analyses and visualisations like those presented in this paper. It contains a handful of user-configurable parameters, which can be used to specify image name metadata, configure filters for the final data table, and specify whether or not to incorporate fluorescence quantification data.

**MDR and CoA** Importantly, the RMarkdown file computes values for the MDR and CoA metrics. As described in the main text, they have been adapted from Zheng et al. 2022[15] to operate on segmented data (rather than raw fluorescence data). In the process, we have made some changes to the algorithms used to calculate these metrics, which we describe below.

MDR describes an organelle's distribution between the nucleus and the cell periphery. We calculate the MDR by applying the following formula to all pixels:

$$\frac{distance\ to\ nucleus}{distance\ to\ nucleus\ +\ distance\ to\ periphery}$$

where these values reflect the Euclidean distances to the nearest pixel of nucleus or cell periphery. We then average this across all pixels within an organelle mask to give the final MDR for the organelle. By this definition, the MDR of the nucleus itself is always zero, although there is an option in the code to instead calculate MDR based on distance to the nucleus *centroid* (or cell centroid, if no nucleus mask is provided), rather than any nucleus pixel.

CoA instead reflects an organelle's asymmetry around the cell. To calculate this value, we divide the cell into an even number of wedges with a fixed angle (5° by default, although this is configurable) around the nucleus centroid. Moving through opposing pairs of wedges, we subtract the number of organelle pixels in each, and tally up the number of residual (i.e. imbalanced) pixels. This number is then normalised to the total number of pixels in the organelle mask to give the CoA.

**Additional metrics** The RMarkdown code additionally calculates a number of metrics related to the distances between different combinations of organelles and contacts (e.g. the average distance from an ER-mitochondrial contact to the nearest lysosome). Because of the very large number of possible combinations, and therefore very large number of associated metrics, we have not included these in our standard analyses, but they are available for targeted interrogation into organelles or contacts of interest.

#### Image presentation

The following lookup tables (LUTs) were used to display the composite images in this paper. All LUTs were accessed via the "NeuroCyto LUTs" update site for ImageJ, by Cristophe Leterrier.

| Organelle | LUT | Source |
|---|---|---|
| Golgi | BIOP Amber | Romain Guiet, EPFL BIOP |
| peroxisomes | BIOP Chartreuse | Romain Guiet, EPFL BIOP |
| ER | CB Bluish Green | Bruno C. Vellutini, after Okabe & Ito |
| mitochondria | BIOP Electric Indigo | Romain Guiet, EPFL BIOP |
| lysosomes | BIOP Bright Pink | Romain Guiet, EPFL BIOP |
| nucleus | Blue | ImageJ default |
| Additional markers | Grays | ImageJ default |

Supplementary Table 1: Lookup tables
